# Perivascular cells support folliculogenesis in the developing ovary

**DOI:** 10.1101/2021.04.26.441453

**Authors:** Shuyun Li, Bidur Bhandary, Tony DeFalco

## Abstract

Granulosa cells, supporting cells of the ovary, are essential for ovarian differentiation by providing a nurturing environment for oogenesis. Sufficient numbers of granulosa cells are vital for establishment of follicles and the oocyte reserve; therefore, identifying the cellular source from which granulosa cells are derived is critical for understanding basic ovarian biology. One cell type that has received little attention in this field is the perivascular cell. Here we use lineage tracing and organ culture techniques in mice to identify ovarian Nestin+ perivascular cells as multipotent progenitors that contribute to granulosa, thecal, and pericyte lineages. Maintenance of these progenitors was dependent on vascular-mesenchymal Notch signaling. Ablation of postnatal Nestin+ cells resulted in a disruption of granulosa cell specification and an increased incidence of polyovular ovarian follicles, thus uncovering key roles for vasculature in ovarian differentiation. These findings may provide new insights into the origins of female gonad dysgenesis and infertility.

## Introduction

Testis and ovary organogenesis are unique processes in which two distinct organs can arise from a common gonadal primordium^1^. Sex determination is initiated around embryonic day (E) 11.5 (E11.5) in mice, at which time testes and ovaries undergo sex-specific molecular and morphogenetic changes^1, 2^. In the absence of the *Sry* gene, the bipotential gonad undergoes ovarian development, ultimately leading to the formation of follicles consisting of a single oocyte surrounded by somatic granulosa cells. The fetal ovary is composed of ovarian surface epithelial cells, endothelial cells forming blood vessels, mesenchymal or stromal cells, granulosa (or pre-granulosa) cells, and germ cells^3^.

Granulosa cells play a critical role in supporting oocyte development by producing multiple hormones and growth factors in immature and mature follicles^4^. The embryonic origins of granulosa cells are still not entirely clear. There are two classes of granulosa precursors that have been described in the fetal mouse ovary. The first class of granulosa cells, which express FOXL2 at early fetal stages, gives rise to medullary primordial follicles that are synchronously activated and become the first wave of active follicles after birth. The second class of follicle cells is composed of LGR5-positive somatic cells, or coelomic epithelial cells at fetal stages, and gives rise to primordial follicles of the adult ovarian cortex that are gradually activated over the reproductive lifespan and, therefore, constitute the definitive oocyte reserve. Both medullary and cortical granulosa cells in the postnatal ovary are marked by the expression of FOXL2^5,6,7^.

Blood vessels are well known for delivering oxygen and nutrients to growing organs. However, recent studies have demonstrated that blood vessels are a critical component of tissue stem cell niches and play essential roles during organogenesis^8,9,10^. Vascular development during gonadogenesis in mice first occurs around E11.5 in both sexes^11^. The vasculature is remodeled to generate a coelomic arterial network in mouse fetal testicular development^11^, and blood vessels are critical for multiple aspects of testis morphogenesis^12, 13^ and for testis differentiation^14^.

In contrast to the fetal testis, little is known about the role of the vasculature during ovary organogenesis^15^. In the mouse fetal ovary, the development of blood vessels does not generate a coelomic arterial vessel. Instead, there are dense medullary networks of vessels, whose function is unclear, near clusters of germ cells known as ovigerous cords or germline cysts^16^. The VEGF (vascular endothelial growth factor) signaling pathway is involved in early follicle activation, as well as angiogenesis, in the postnatal rat ovary^17^; however, the factors regulating ovary patterning, especially in the interstitial compartment during fetal and perinatal stages, remain largely unaddressed. In particular, ovarian perivascular cells are poorly understood, but in light of a recent single-cell RNA-Seq study indicating that perivascular cells are a major constituent of human ovaries^18^, it is likely that perivascular cells play unappreciated roles in ovarian development. As of yet, the cell types that arise from perivascular cells in the ovary have not been well-defined.

Notch signaling integrates niche signals to direct cell differentiation during organ morphogenesis, including testis and ovary organogenesis^14, 19, 20^. Activation of the Notch pathway in the fetal and postnatal ovary has been implicated in vascular development, preovulatory granulosa cell differentiation, and regulation of meiosis in oocytes^19, 21, 22^. However, the cellular players involved in fetal ovarian Notch signaling are poorly understood and, in particular, links between Notch signaling, vascular-mesenchymal interactions, and perivascular cells remain major outstanding questions in the field.

In this study, we have used genetic lineage tracing assays and *ex vivo* organ culture approaches to address the origins and roles of vasculature and perivascular cells in the mouse ovary. We show that ovarian vasculature is critical for maintaining a multipotent progenitor niche that gives rise to several cell types, such as granulosa cells, theca cells, and pericytes. These perivascular progenitors express Nestin, a marker for neural stem cells and stem-like progenitor cells in various organs^14, 23,24,25^. Inhibition of vasculature disrupted the maintenance of Nestin+ progenitor cells and induced their precocious differentiation into granulosa cells during a specific developmental window. Maintenance of these progenitors was dependent on vascular-dependent Notch signaling, demonstrating that Notch signaling provides a controlled environment for maintaining a balance between perivascular progenitor cells and differentiated granulosa cells in the perinatal ovary. Finally, we specifically ablated Nestin+ cells and found that granulosa cell specification was disrupted, leading to defective folliculogenesis and an increased occurrence of polyovular follicles. Our results uncover a new instructive role for ovarian vasculature in the progenitor niche for granulosa cells, which is essential for establishing follicles, the structural and functional units of the ovary. These findings provide new insights into basic mechanisms underlying organogenesis and highlight how vasculature and perivascular cells are vital components of ovarian development.

## Results

### Undifferentiated perivascular cells express Nestin in the fetal ovary

Given that testes and ovaries arise from a common primordium, i.e., the bipotential gonad, we investigated whether Nestin+ progenitors exist in the fetal ovary. Immunofluorescence analyses of E12.5 fetal ovaries revealed that Nestin protein was expressed in stromal cells adjacent to blood vessels (Fig. 1a). Immunofluorescence for ICAM2, a vascular-endothelial-cell-specific marker within the gonad, confirmed that Nestin was not expressed in endothelial cells but rather in perivascular mesenchymal cells (Fig. 1a).

**Fig. 1.**
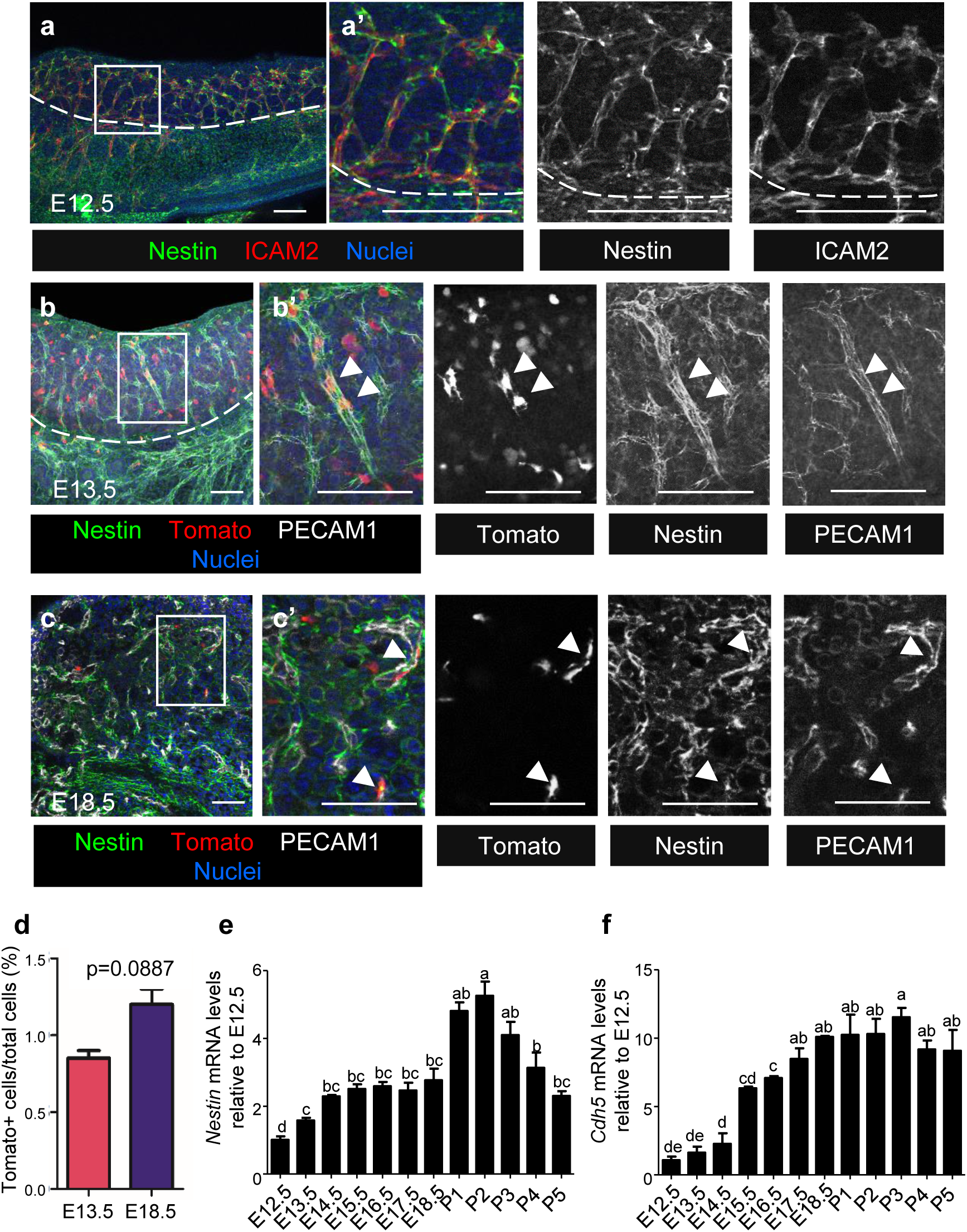
Ovarian Nestin+ cells are spatially and temporally associated with vascular development. (a) Immunofluorescence image of an E12.5 wild-type CD-1 fetal ovary, showing Nestin expression in cells adjacent to ICAM2+ vasculature. (b,c) Images of E13.5 (b) and E18.5 (c) *Nestin*-CreER;*Rosa*-Tomato fetal ovaries exposed to 4-OHT at E12.5, showing Nestin and Tomato expression in perivascular cells (arrowheads). Dashed lines indicate gonad-mesonephros border; a’-c’ are higher-magnification images of the boxed regions in a-c. Scale bars: 50 μm. (d) Flow cytometry analyses of Tomato+ cells in E13.5 and E18.5 *Nestin*-CreER; *Rosa*-Tomato fetal ovaries exposed to 4-OHT at E12.5. (e,f) Whole ovary qRT-PCR analyses of *Nestin* (e) and *Cdh5* (f) mRNA expression at various developmental stages. Letters indicate statistical differences via a two-tailed Student’s t-test (*P*<0.05). qRT-PCR values in e and f are presented as mean ± SD.

*Nestin*-CreER;*Rosa*-Tomato mice were used to permanently label Nestin+ perivascular cells and their progeny in the ovary. To induce CreER activity, we injected 4-hydroxytamoxifen (4-OHT) intraperitoneally into pregnant females at E12.5, upon the first appearance of differentiated pre-granulosa cells. Analyses of E13.5 and E18.5 Nestin-CreER;*Rosa*-Tomato fetal ovaries revealed that Nestin protein and Tomato were specifically enriched in cells adjacent to endothelial cells (Fig. 1b,c). Flow cytometric analyses revealed no significant difference in the percentage of Tomato+ cells within E13.5 versus E18.5 ovaries (Fig. 1d), suggesting Tomato-positive cells initially labeled at E12.5 did not significantly proliferate relative to other cell types. However, the mRNA levels of *Nestin* progressively increased in the ovary during fetal and postnatal stages. Compared with E12.5 ovaries, *Nestin* mRNA levels were significantly increased during later fetal stages and peaked at postnatal (P) day 1-3 (P1-3) (Fig. 1e). *Cdh5* (also called *VE-Cadherin*), an endothelial-specific gene, followed a similar pattern of expression during fetal stages and was maintained after birth (Fig. 1f). These flow cytometric and *Nestin* mRNA expression data suggest that perivascular Nestin+ cells initially labeled at E12.5 did not self-renew or expand, but instead new Nestin+ cells were continuously induced or recruited in the ovary during later fetal and perinatal stages.

### Perivascular Nestin+ cells are derived from gonadal NR5A1+ somatic cells

We next sought to determine the origin of Nestin+ cells, which in the fetal ovary could potentially be the surface epithelium, gonadal mesenchyme, or mesonephros^26^. To test whether Nestin+ cells were derived from ovarian surface epithelium, MitoTracker dye was used to label the surface cells of E11.5 and E12.5 fetal ovaries, followed by *ex vivo* organ culture. At initial timepoints, the dye was clearly confined to the surface epithelium of E11.5 and E12.5 gonads (Supplementary Fig. 1a-f). Nestin+ cells within the interior of the gonad after culture for 48 hours were not labeled with MitoTracker (Supplementary Fig. 1c,d). However, after culture for 48 hours, Mitotracker-labeled epithelial cells that remained on the surface started to express Nestin (Supplementary Fig. 1b,f). These data suggest that gonadal mesenchymal Nestin+ cells adjacent to vasculature did not arise from the coelomic surface epithelium of the fetal ovary.

To determine whether Nestin+ cells are derived from the gonad or mesonephros, we examined the expression of NR5A1 (also called SF1 or Ad4BP), which is restricted to early gonadal somatic cells, but is not expressed in the mesonephros, and gives rise to pre-granulosa cells and potential steroidogenic precursor cells^27, 28^. In E13.5 *Nestin*-CreER;*Rosa*-Tomato ovaries exposed to 4-OHT at E12.5, most Tomato-labeled cells overlapped with NR5A1 (Supplementary Fig. 1g). In contrast, most Tomato-labeled cells did not colocalize with WT1, which is expressed in both gonad and mesonephric mesenchyme and contributes to the theca cell lineage^29^ (Supplementary Fig. 1h). Our data suggest that most ovarian perivascular Nestin+ cells arise from gonadal NR5A1+ somatic cells arising at or prior to E11.5, indicating that these early cells potentially give rise to pre-granulosa cells.

### Nestin+ cells in the developing ovary give rise to granulosa cells and theca cells

To determine the adult ovarian cell types that arise from perivascular Nestin+ cells, we examined P60 *Nestin*-CreER;*Rosa*-Tomato ovaries that were exposed to 4-OHT at E12.5. By P60, Tomato-labeled cells did not express Nestin, indicating that Nestin-derived cells had differentiated and subsequently lost expression of Nestin (Fig. 2a). To address the hypothesis that Nestin+ cells are granulosa cell precursors, we examined the granulosa cell marker FOXL2. A subset of Tomato-labeled cells expressed FOXL2 (Fig. 2b), indicating that Nestin+ cells are progenitors of granulosa cells. However, the number of Tomato+ cells was likely underestimated at P60, because the number of 4-OHT-induced Tomato+ cells at 12.5 was limited (likely due to CreER inefficiency) and new Nestin+ progenitors likely emerged following the continued formation of blood vessels during development. To further examine this hypothesis, we administered 4-OHT at E15.5, E18.5, P2, or P4 in *Nestin*-CreER;*Rosa*-Tomato mice to lineage-trace Nestin+ cells more efficiently (based on ovarian *Nestin* mRNA levels; see Fig. 1e). After 24 hours, Nestin+ cells were labeled with Tomato. Spindle-shaped Tomato-positive cells were adjacent to blood vessels and did not co-express FOXL2 at E16.5 (Supplementary Fig. 2a). At P1, P3, and P5, occasional round-shaped Tomato-labeled co-expressed FOXL2, but the vast majority of initially labeled Tomato+ cells were spindle-shaped and were specifically localized adjacent to vasculature (Supplementary Fig. 2b-d).

**Fig. 2.**
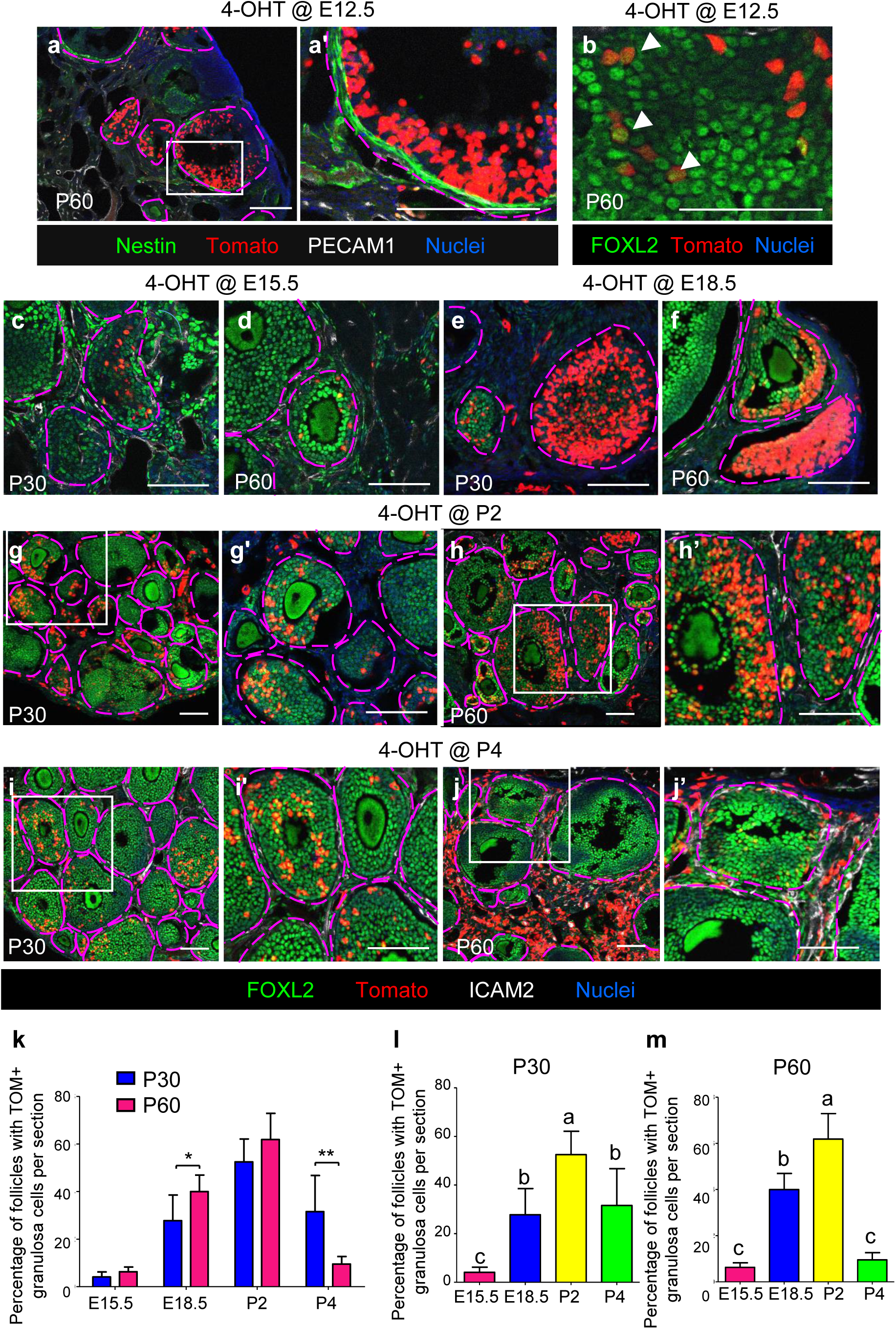
Fetal and postnatal perivascular Nestin+ cells give rise to granulosa cells. (a-j) Long-term lineage tracing of fetal and postnatal Nestin+ cells in P30 (c,e,g,i) and P60 (a,b,d,f,h,j) *Nestin*-CreER;*Rosa*-Tomato ovaries exposed to 4-OHT at E12.5 (a,b), E15.5 (c,d), E18.5 (e,f), P2 (g,h), or P4 (i,j). (a) Differentiated Tomato-labeled cells did not express Nestin. (b) A subset of Tomato-labeled cells co-expressed the granulosa cell marker FOXL2 (arrowheads). (c,d) When exposed to 4-OHT at E15.5, few follicles contained Tomato-labeled granulosa cells. (e-j) When exposed to 4-OHT at E18.5, P2, or P4, numerous follicles with Tomato-labeled granulosa cells were observed. (i,j) When exposed to 4-OHT at P4, most Tomato-labeled cells at P60 were located in the interstitium, with few granulosa cells within each follicle labeled with Tomato. a’ and g’-j’ are higher-magnification images of the boxed regions in a and g-j. Scale bars: 50 μm. (k-m) Graph shows percentage of primary, secondary, and antral follicles containing Tomato+ granulosa cells in P30 and P60 *Nestin*-CreER;*Rosa*-Tomato ovaries exposed to 4-OHT at E15.5, E18.5, P2, or P4. Data are presented as mean ± SD.

Similar to E12.5 injection, ovaries exposed to 4-OHT at E15.5 showed only 1-2 follicles in each section containing Tomato-labeled FOXL2+ cortical granulosa cells at P30 and P60 (Fig. 2c,d,k). Additionally, we observed only a few Tomato-positive cells that expressed HSD3B1, a marker of thecal and interstitial steroidogenic cells in the adult ovarian stroma^30^, in P30 and P60 samples (Supplementary Fig. 3a,b). We also saw occasional Tomato+ vascular smooth muscle cells (VSMCs) expressing alpha-smooth muscle actin (ACTA2; also called α-SMA) and pericytes expressing CSPG4 (also called NG2) in the theca layer of follicles^31,32,33,34^ in P30 and P60 ovaries that were exposed to 4-OHT at E15.5 (Supplementary Fig. 3c-f).

However, after administering 4-OHT to E18.5 ovaries, which mainly contained germline cysts at that stage, Tomato+ cells were observed in a larger subset of granulosa cells of cortical follicles (Fig. 2e,f). According to follicle quantification data, progeny of Nestin+ cells in E18.5 fetal ovaries were biased towards granulosa cells in secondary-wave follicles (i.e., observed more often at P60 versus P30) (Fig. 2k). Only a few Tomato+ cells expressed HSD3B1 at P30 (Supplementary Fig. 3g), whereas a larger number of Tomato+ cells colocalized with HSD3B1 at P60 (Supplementary Fig. 3h). In P30 and P60 ovaries, many Tomato+ cells were observed in the theca layer, expressing ACTA2 or CSPG4 (Supplementary Fig. 3i-l).

After administering 4-OHT at P2 (when germline cysts break apart and individual oocytes become surrounded by somatic cells to form primordial follicles), more Tomato+ cells overlapped with FOXL2 as compared to E18.5 injection (Fig. 2g,h,k). Additionally, Tomato+ cells co-expressed CSPG4 and ACTA2 (markers of theca cells; Supplementary Fig. 3o-r), as well as HSD3B1 (a marker of steroidogenic cells throughout the ovarian interstitium; Supplementary Fig. 3m,n).

When pups were administered 4-OHT at P4, fewer Tomato+ cells were FOXL2+ granulosa cells in adult ovaries, while many Tomato+ cells were interstitial mesenchymal cells (Fig. 2i,j). Nestin+ cells in P4 ovaries mainly contributed to granulosa cells in medullary first-wave follicles (Fig. 2k). Few Tomato+ cells expressed HSD3B1 at P30; more Tomato+ cells expressed HSD3B1 in P60 ovaries as compared to P2 injection (Supplementary Fig. 3s,t). Likewise, Tomato+ cells were observed in the theca layer (Supplementary Fig. 3u-x).

The percentage of follicles with Tomato+ granulosa cells in *Nestin*-CreER;*Rosa*-Tomato ovaries (exposed to 4-OHT at different stages) showed that P2-derived Nestin+ cells contributed to granulosa cells in both first- and second-wave follicles (Fig. 2l,m), whereas P4-derived cells were less likely to give rise to granulosa cells in second-wave follicles and more likely to give rise to interstitial cell types such as steroidogenic cells in P60 ovaries. These data provide evidence that fetal and early postnatal Nestin+ cells can give rise to granulosa cells, as well as theca cells and interstitial stromal cells, in a stage-specific manner.

### Maintenance of Nestin-expressing cells requires blood vessels

To explore the requirement of vasculature for Nestin expression in fetal and postnatal ovaries, we disrupted vasculature *ex vivo* with a VEGF signaling blocker (VEGF receptor inhibitor VEGFR-TKI II). VEGFR-TKI II treatment significantly reduced expression of the endothelial-specific markers ICAM2 and *Cdh5* (Fig. 3a,b), demonstrating the effectiveness of vascular inhibition. We also observed a reduction in mRNA and protein levels of *Nestin*, indicating that ovarian *Nestin* expression is dependent on the presence of blood vessels (Fig. 3a,c).

**Fig. 3.**
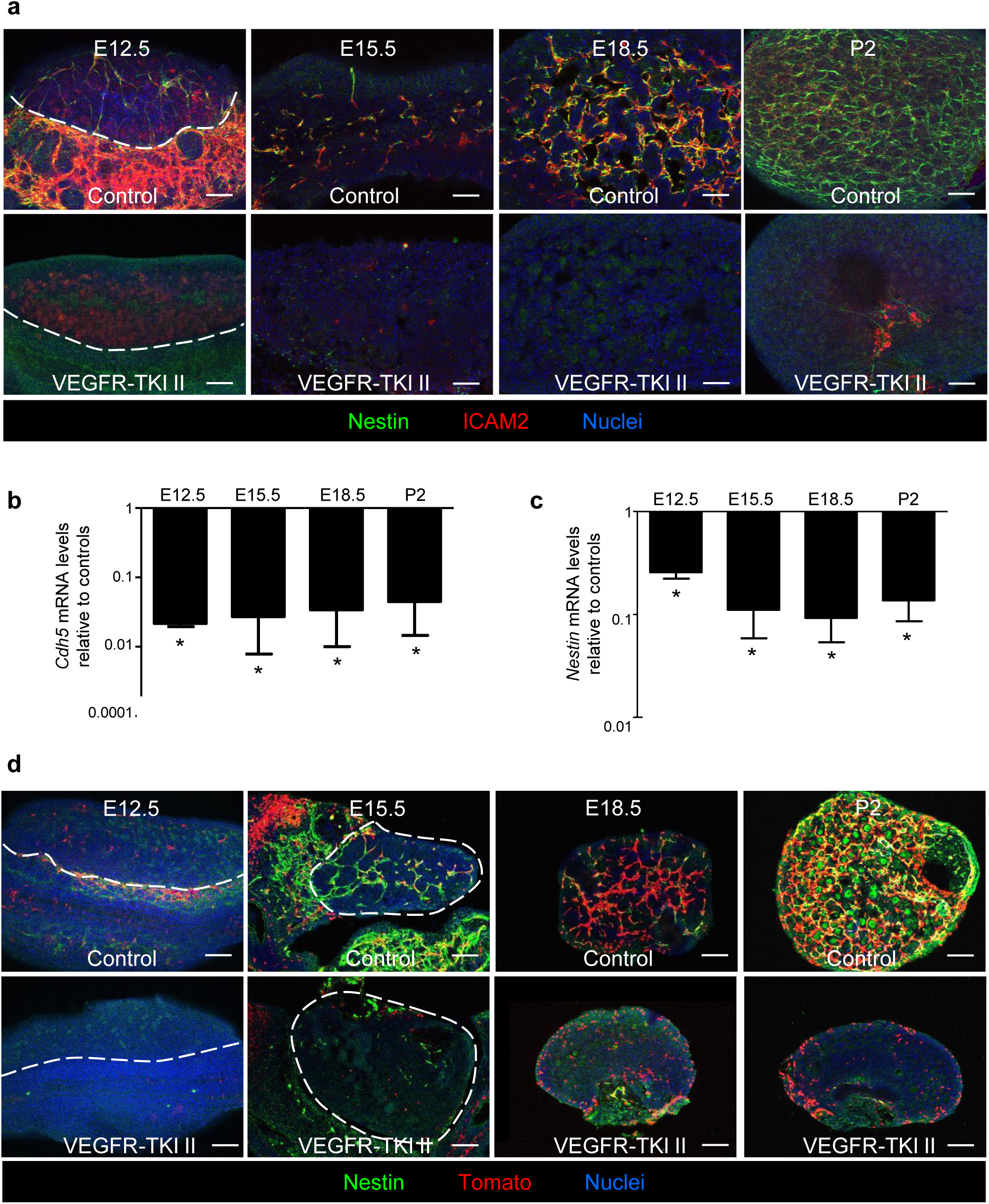
Vasculature is required for maintenance of perivascular Nestin+ cells. Immunofluorescence images (a,d) and qRT-PCR analyses (b,c) of E12.5, E15.5, E18.5, or P2 wild-type CD-1 ovaries cultured *ex vivo* with DMSO (control) or VEGFR-TKI II (VEGFR inhibitor). (a) Depletion of vasculature resulted in decreased expression of Nestin and ICAM2. (b,c) qRT-PCR analyses showing fold change in *Nestin* (b) and *Cdh5* (c) mRNA expression in whole fetal and postnatal ovaries after vascular disruption. \**P*<0.05; two-tailed Student’s t-test. qRT-PCR values in b and c are presented as mean ± SD. (d) *Ex vivo* vascular disruption of *Nestin*-CreER;*Rosa*-Tomato fetal and postnatal ovaries, showing that Tomato-positive cells were mostly spindle-shaped in control ovaries but were round-shaped in vascular-depleted ovaries. Dashed lines indicate gonad-mesonephros border. Scale bars: 50 μm.

To directly visualize whether Nestin-derived cells remained in the ovary in the absence of vasculature, we disrupted vasculature in *Nestin*-CreER;*Rosa*-Tomato fetal and postnatal ovaries (Fig. 3d). Tomato+ cells were no longer detectable after disruption of vasculature at E12.5. In E15.5 fetal ovaries, some Tomato-labeled cells, which had a different cellular morphology as compared to controls, remained in vascular-disrupted gonads. Most spindle-shaped Tomato-positive cells in control ovaries were next to blood vessels in the medullary region; however, after vascular disruption at E18.5, Tomato-positive cells changed from spindle-shaped into round-shaped, and they were enriched in the ovarian cortex. However, ablation of vasculature in P2 ovaries only resulted in the reduction of perivascular Tomato-labeled cells but did not increase the number of Tomato+ cortical cells (Fig. 3d).

### Vasculature is required to block premature granulosa cell differentiation

We next used *Nestin*-CreER;*Rosa*-Tomato ovaries to examine the specific fate of Nestin-derived cells after vascular disruption. As expected, VEGFR-TKI II significantly decreased ICAM2 expression (Fig. 4a-c). Whereas E15.5 and E18.5 control ovaries contained Tomato+ spindle-shaped perivascular cells, Tomato+ cells that remained in E15.5 and E18.5 vascular-disrupted ovaries were cortical FOXL2+ cells (Fig. 4a,b). At P2, vascular depletion only decreased medullary Tomato+ cells, whereas cortical differentiated Tomato+ cells (FOXL2+) were unaffected (Fig. 4c). These data indicated that the disruption of vasculature induced differentiation of perivascular Nestin+ cells into FOXL2+ pre-granulosa cells most notably at E18.5. To further test this hypothesis, *Foxl2* expression was examined in VEGFR-TKI II-treated ovaries as compared to control ovaries. Blocking vasculature did not significantly affect *Foxl2* mRNA levels in E12.5, E15.5, and P2 ovaries, but did significantly increase *Foxl2* mRNA levels (∼6-fold increase) in E18.5 fetal ovaries (Fig. 4d). Taken together, these data suggest that Nestin+ progenitors prematurely or ectopically differentiated into pre-granulosa cells in the absence of vasculature during a specific time window.

**Fig. 4.**
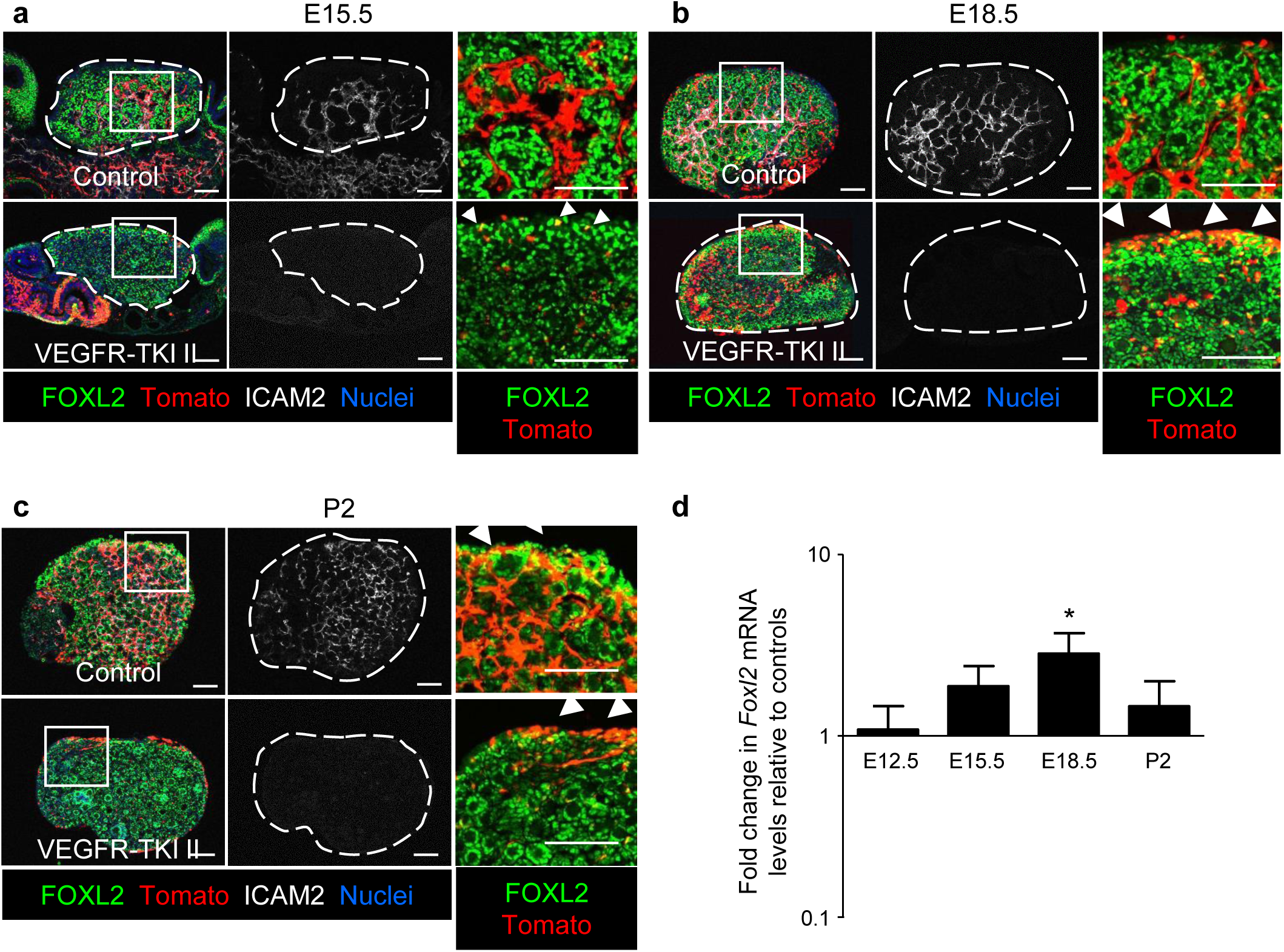
Vascular disruption results in differentiation of Nestin+ cells into cortical pre-granulosa cells. Immunofluorescence images (a-c) and qRT-PCR analyses (d) of E15.5 (a), E18.5 (b), or P2 (c) *Nestin*-CreER;*Rosa*-Tomato ovaries cultured *ex vivo* with DMSO (control) or VEGFR-TKI II. (a) Vascular disruption at E15.5 resulted in reduced numbers of Tomato+ and ICAM2+ cells. (a) After depletion of vasculature at E18.5, spindle-shaped Tomato-labeled cells in the medulla were decreased, while Tomato+ cells, co-expressing FOXL2, were enriched in the ovarian cortex. (c) After vascular disruption at P2, medullary spindle-shaped cells labeled with Tomato were decreased, with no change in cortical Tomato-labeled cells. Arrowheads throughout indicate cortical FOXL2+ cells. Dashed outlines in a-c indicate gonad boundary. Images on the right in each panel are higher-magnification images of the boxed regions in the left image. Scale bars: 50 μm. (d) qRT-PCR analyses showing fold change in *Foxl2* mRNA levels in fetal and postnatal ovaries after vascular disruption relative to controls. \**P*<0.05; two-tailed Student’s t-test. qRT-PCR values in d are presented as mean ± SD.

### Active Notch signaling in female gonads is dependent on vasculature

Active Notch signaling within fetal ovaries is restricted to somatic cells^35^; however, it is unknown whether Notch signaling acts as the bridge between vasculature and somatic cells in fetal ovaries. Therefore, to investigate Notch signaling activity in fetal and postnatal ovaries, we performed immunofluorescence analyses using a *CBF*:H2B-Venus line as a Notch signaling reporter^36^. In E15.5 female gonads, little Notch activity was observed in Nestin+ cells; in contrast, more perivascular Nestin+ cells expressed Venus at E18.5 and P2 (Supplementary Fig. 4a). In P4 ovaries, Notch activity was infrequently observed in perivascular Nestin+ somatic cells (Supplementary Fig. 4a). Relative to E12.5 ovaries, mRNA levels of the Notch target genes *Hes1, Heyl, Hey1* were up-regulated at E18.5 and peaked at P1, whereas mRNA levels of *Hes5* did not change during E18.5 to P4 (Supplementary Fig. 4b-e). Taken together, Notch signaling was most active in the ovary during E18.5 to P4. To determine if pre-granulosa cells underwent active Notch signaling, we examined FOXL2 expression in P2 *CBF*:H2B-Venus ovaries. Immunofluorescence analyses showed that perivascular Venus-positive cells did not express FOXL2; however, some Venus-positive cells in the ovarian cortex that were not adjacent to vasculature co-expressed FOXL2 (Supplementary Fig. 4f, white arrowheads).

To further analyze blood vessel control of Notch activity, we disrupted vasculature *ex vivo* in *CBF*:H2B-Venus fetal and postnatal ovaries. In E12.5 and E15.5 fetal ovaries, Venus expression was absent in gonads after vascular disruption; however, some Venus remained in the mesonephros (Fig. 5a,b). At E18.5, blockade of vasculature resulted in a reduction of cortical Venus and absence of medullary Venus expression (Fig. 5c). However, vascular blocking in P2 ovaries only disrupted active Notch signaling in the medulla and did not affect Notch activity in the cortex (Fig. 5d). Next, we examined the mRNA expression of Notch target genes to further verify the role of the vasculature in driving active Notch signaling. *Hes1*, *Hes5*, *Hey1*, and *Heyl* mRNA levels were all significantly decreased after disruption of vasculature at E12.5, E15.5, and E18.5, but not at P2 (Fig. 5e).

**Fig. 5.**
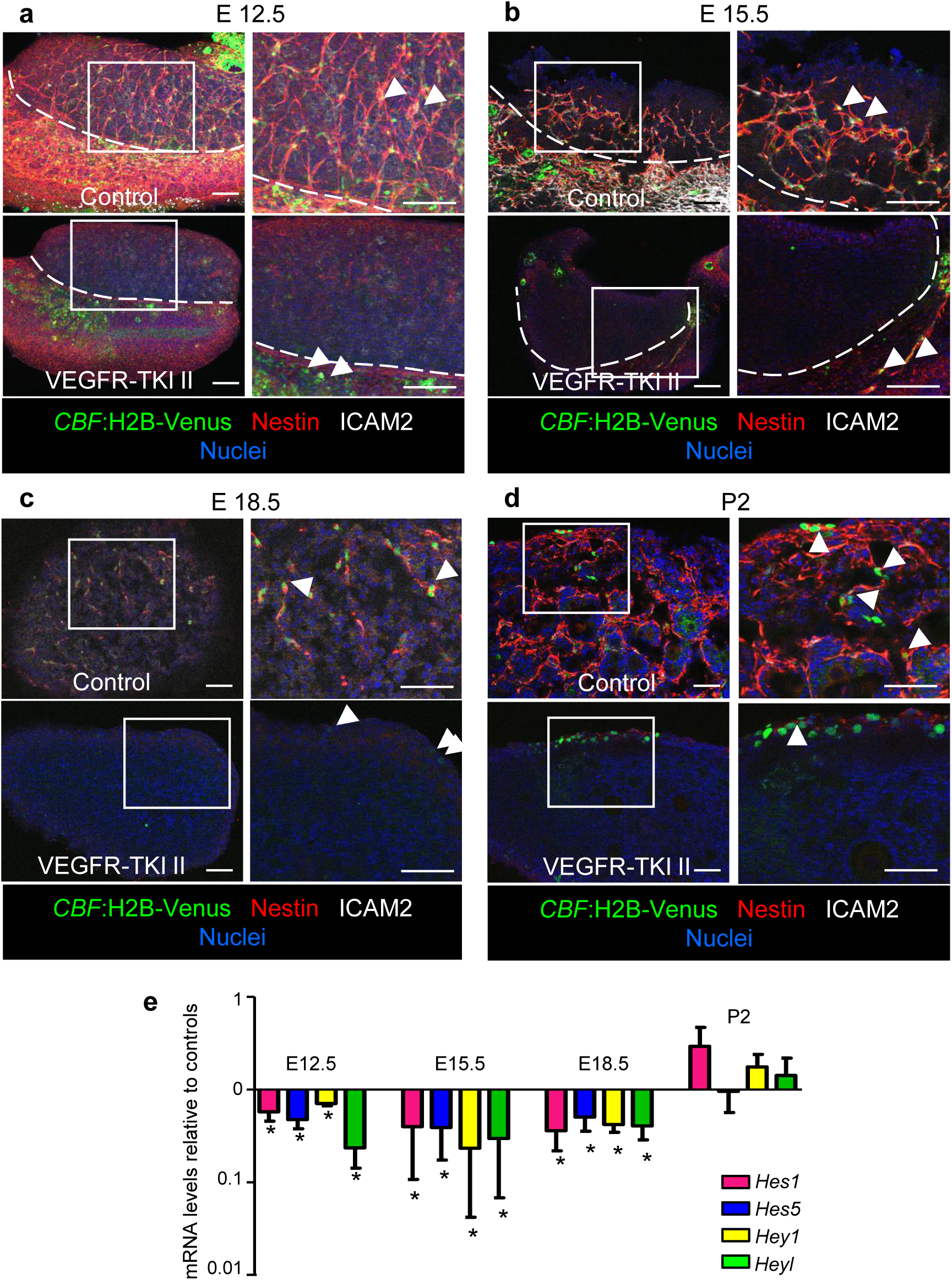
Vasculature is essential for medullary Notch activity in fetal and postnatal ovaries. Immunofluorescence images (a-d) and qRT-PCR analyses (e) of E12.5 (a), E15.5 (b), E18.5 (c) or P2 (d) *CBF*:H2B-Venus ovaries (Notch signaling reporter) cultured *ex vivo* with DMSO (control) or VEGFR-TKI II. At E12.5 (a) and E15.5 (b), depletion of vasculature reduced Notch signaling (Venus expression) in the gonad but not in the mesonephros (arrowheads in a and b). In E18.5 (c) and P2 (d) ovaries, vascular disruption decreased medullary Notch activity; however, Venus+ cells were detected on the gonad surface (arrowheads in c and d). Images on the right in each panel are higher-magnification images of the boxed regions in the left image. Dashed lines in a and b indicate gonad-mesonephros border. Scale bars: 50 μm. (e) qRT-PCR analyses showing fold change in Notch target gene (*Hes1*, *Hes5*, *Hey1*, *Heyl*) mRNA levels after disruption of vasculature at E12.5, E15.5, E18.5, or P2. \**P*<0.05; two-tailed Student’s t-test. qRT-PCR values in e are presented as mean ± SD.

### Notch signaling regulates differentiation of ovarian Nestin+ progenitors

Whether Notch signaling is a critical aspect of vascular-mesenchymal interactions during ovarian differentiation in this context has not been definitively assessed. Specifically, we hypothesized that Notch signaling is responsible for the maintenance of ovarian perivascular Nestin+ cells; thus, we blocked Notch activity by using the γ-secretase inhibitor DAPT in fetal and postnatal ovaries *ex vivo*. Expression of Notch target genes was significantly reduced in DAPT-treated groups, demonstrating that Notch signaling was effectively blocked (Fig. 6a). Blockade of Notch activity reduced Nestin protein expression in ovaries at E15.5, E18.5, and P2 (Fig. 6b-d). However, DAPT also decreased ICAM2 expression at E18.5 and P2 (Fig. 6c,d,f), indicating a disruption in vasculature. We next examined *Nestin* and *Cdh5* mRNA expression in DAPT-treated ovaries. *Nestin* mRNA levels were significantly down-regulated by DAPT at E15.5, E18.5, and P2 (Fig. 6e). Consistent with immunofluorescence images, DAPT only affected *Cdh5* mRNA expression at E18.5 and P2 but not at E15.5 (Fig. 6f), indicating there is a stage-specific effect of Notch signaling upon ovarian vascular development. To determine whether the reduction of *Nestin* mRNA expression in DAPT-treated ovaries was due to down-regulation of *Cdh5* mRNA expression (i.e., loss of vasculature), we treated E18.5 ovaries with DAPT for 12, 18, or 24 hours. DAPT specifically decreased *Nestin* mRNA levels when cultured for 18 hours (i.e., did not significantly affect *Cdh5* levels) (Supplementary Fig. 5a), suggesting that reduction of *Nestin* expression is not solely due to a reduction in vasculature after DAPT treatment.

**Fig. 6.**
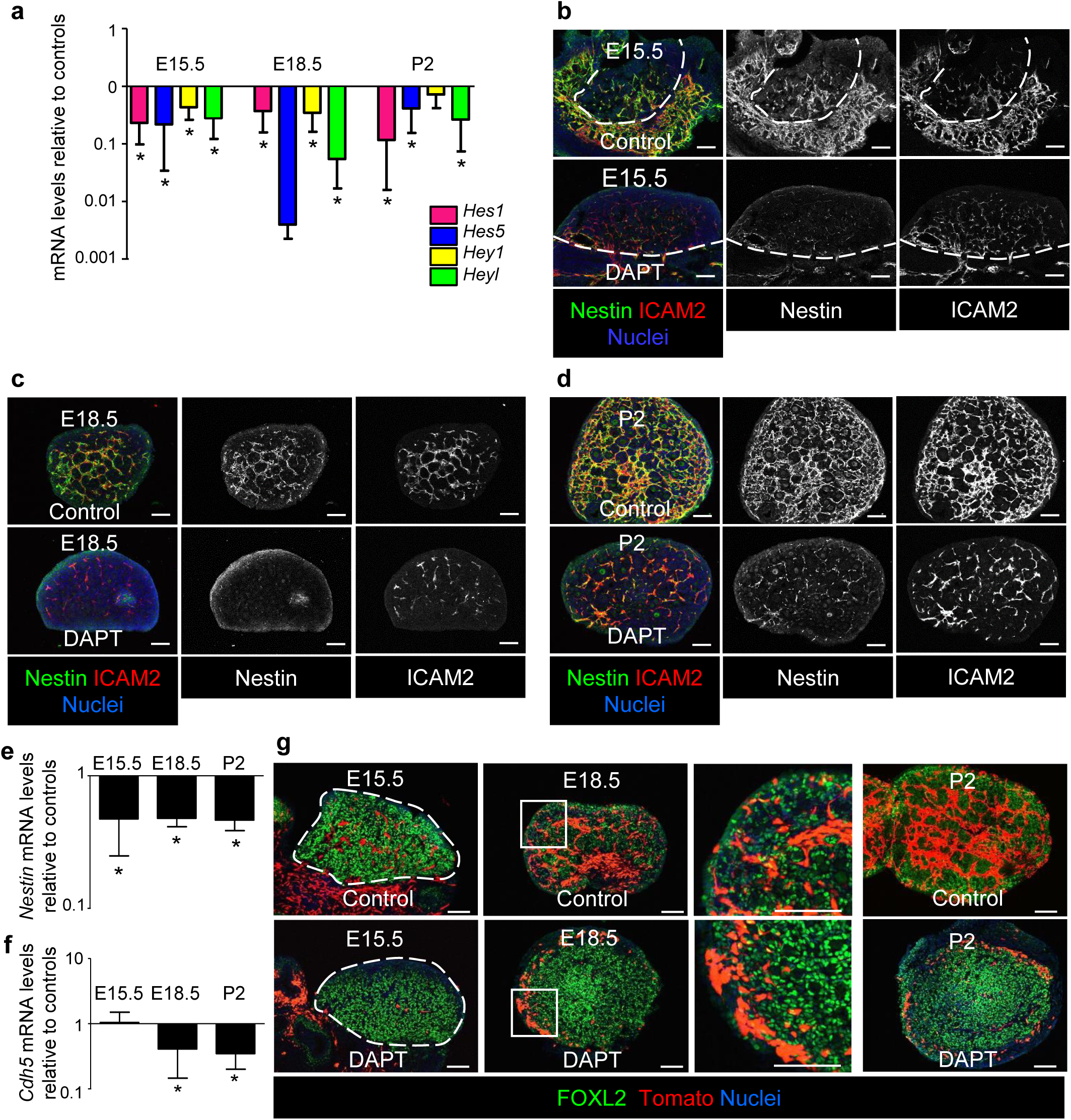
Disruption of Notch signaling induces differentiation of Nestin-derived cells into pre-granulosa cells. (a-f) Immunofluorescence images (b-d) and qRT-PCR analyses (a,e,f) of E15.5 (b), E18.5 (c), or P2 (d) wild-type CD-1 ovaries cultured *ex vivo* with DMSO (control) or DAPT (Notch signaling inhibitor). (a) qRT-PCR analyses showing fold change in Notch target gene (*Hes1*, *Hes5*, *Hey1*, *Heyl*) mRNA levels after blockade of Notch signaling via DAPT at E15.5, E18.5, or P2. (b-d) After disruption of Notch signaling at E15.5 (b), only Nestin expression decreased, whereas at E18.5 (c) and P2 (d) DAPT treatment decreased both Nestin and ICAM2 expression. (e,f) qRT-PCR analyses showing fold change in *Nestin* (e) and *Cdh5* (f) mRNA levels in ovaries cultured with DAPT at E15.5, E18.5, or P2. (g) *Ex vivo* culture of E15.5, E18.5, or P2 *Nestin*-CreER;*Rosa*-Tomato fetal and postnatal ovaries. At E15.5, DAPT treatment reduced Tomato+ cells, which did not express FOXL2, but at E18.5 DAPT induced round-shaped cortical Tomato+ cells and reduced medullary spindle-shaped Tomato+ cells. At P2, DAPT only decreased medullary Tomato+ cells with no impact on cortical Tomato+ cells. Dashed lines throughout indicate gonad outlines. Scale bars: 50 μm. *, *P*<0.05; two-tailed Student’s t-test. qRT-PCR values in a, e, and f are presented as mean ± SD.

To further verify that Notch signaling is a regulator of Nestin+ cells, *Nestin* CreER;*Rosa-*Tomato;*Rosa*-NICD embryos were exposed to 4-OHT at E18.5 to stimulate Notch signaling through CreER-mediated, ligand-independent expression of the Notch intracellular domain (NICD). Perivascular Tomato+ cells were induced in NICD+ ovaries (Supplementary Fig. 5b,c). We used Ki67 and phospho-histone H3 (pHH3) to determine cell cycle status and proliferation of Tomato+ cells, respectively. However, Tomato+ cells did not express MKI67 or phospho-Histone H3 (Supplementary Fig. 5b,c), indicating these cells did not proliferate from already existing Tomato+ cells. These results suggest that active Notch signaling regulates Nestin expression.

To examine the fate of Nestin+ cells after blockade of Notch signaling, we inhibited Notch activity in *Nestin*-CreER;*Rosa*-Tomato ovaries via DAPT treatment *ex vivo* (Fig. 6g). In E15.5 ovaries, DAPT treatment reduced the number of Tomato+ cells, which did not express FOXL2. Similar to vascular disruption, at E18.5 medullary perivascular Tomato+ cells were decreased and were now enriched in the ovarian cortex. At P2, Notch blockade only reduced medullary perivascular Tomato+ cells but did not affect cortical Tomato+ cells, which had differentiated into pre-granulosa cells (Fig. 6g). Our data suggest that blockade of Notch signaling induced the differentiation of Nestin+ cells into cortical pre-granulosa cells and reduced medullary perivascular undifferentiated Nestin+ cells at E18.5. Furthermore, while Notch activity depended on vasculature in earlier fetal stages, active Notch signaling might contribute to vascular formation in later stages.

### Nestin+ cells are required for folliculogenesis

If perivascular Nestin+ cells represent pre-granulosa progenitors, we anticipated that ablating these cells would compromise granulosa cell differentiation or follicular development during ovary organogenesis. To specifically deplete Nestin+ progenitors from the ovary, we used a Cre-responsive *Rosa*-eGFP-DTA mouse line^37^ driven by *Nestin*-CreER in which diphtheria toxin is only expressed in CreER-active cells, thus inducing apoptosis only in perivascular cells. We injected 4-OHT to deplete Nestin+ cells at P2 and P4 (Fig. 7a), at the stages with highest ovarian *Nestin* expression. Compared with *Nestin*-CreER;*Rosa*-Tomato controls lacking *Rosa-*eGFP-DTA, Tomato expression was absent in P7 and P21 *Nestin*-CreER;*Rosa*-Tomato;*Rosa-*eGFP-DTA (DTA+) ovaries, indicating efficient ablation of Nestin+ cells (Supplementary Fig. 6).

**Fig. 7.**
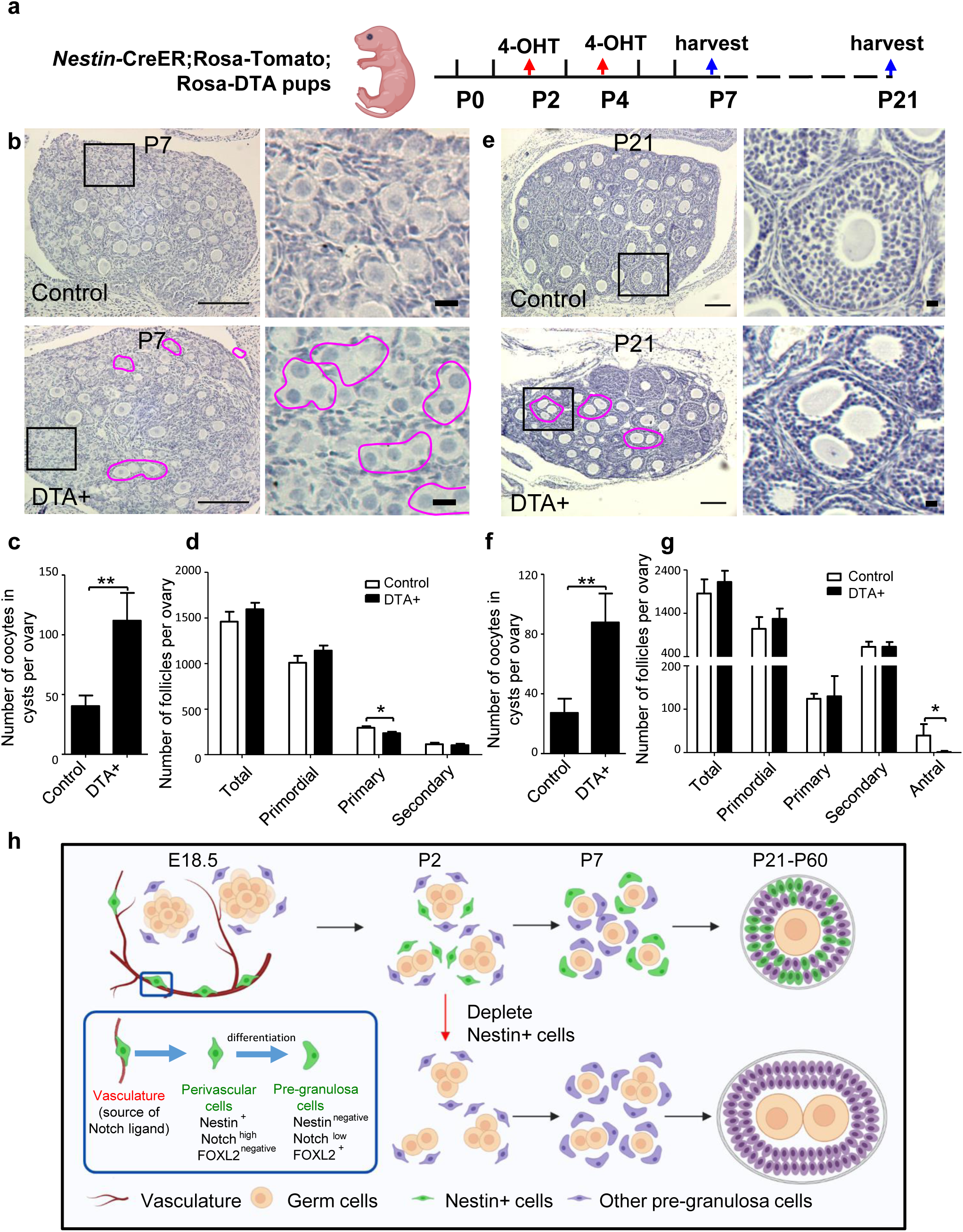
Postnatal Nestin-derived cells are required for folliculogenesis. (a) Cartoon depicting experimental strategy for ablating postnatal Nestin+ cells in *Nestin*-CreER; *Rosa*-Tomato; *Rosa*-eGFP-DTA postnatal ovaries via exposure to 4-OHT at P2 and P4. (b,c) Hematoxylin staining (b) and follicle quantification (c) revealed that cyst breakdown was disrupted in Nestin-cell-depleted (DTA+) P7 ovaries (magenta outlines), whereas primordial follicles (right panels in b) assembled normally in controls. (d) Quantification of follicle types in P7 control and Nestin-cell-depleted (DTA+) ovaries. (e,f) Hematoxylin staining (e) and follicle quantification (f) revealed an increase in remnant germline cysts (i.e., polyovular follicles; magenta outlines) at P21 after depletion of Nestin+ cells. Images on the right in b and e are higher-magnification images of the boxed regions in the left image. (g) Quantification of follicle types in P21 control and Nestin-cell-depleted (DTA+) ovaries, showing lack of antral follicle formation in DTA+ ovaries. *, *P*<0.05, **, *P*<0.01; two-tailed Student’s t-test. Values in c-g are presented as mean ± SD. Thin scale bars: 50 μm; thick scale bars: 5 μm. (h) Cartoon model depicting the dynamic expression pattern and function of Notch-signaling-regulated perivascular Nestin+ cells in the ovary. Cartoon was created using BioRender (biorender.com).

We then assessed how the loss of Nestin+ perivascular cells affected ovarian morphology and histology. At P7, when control follicles normally contain single oocytes, in DTA+ ovaries some abnormal oocyte cysts containing multiple germ cells (i.e., polyovular follicles) remained in the medulla, and more oocyte cysts were observed in the cortex (Fig. 7b,c), indicative of defective oocyte cyst breakdown. We next classified and counted follicles (primordial, primary, secondary, and antral follicles) according to established morphological criteria^38^. Total follicle number in P7 DTA+ ovaries was similar compared to control ovaries; however, Nestin-cell-depleted ovaries exhibited a significantly reduced number of primary follicles, while numbers of primordial and secondary follicles were unchanged at P7 (Fig. 7d).

While control P21 ovaries exhibited normal, ongoing follicular progression, DTA+ ovaries contained numerous polyovular ovarian follicles and exhibited defects in folliculogenesis, showing a significantly reduced number of antral follicles as compared to controls (Fig. 7e-g). Some polyovular follicles were observed in P21 control ovaries, which might be an effect of postnatal 4-OHT injection, but at a rate significantly lower than DTA+ ovaries. (Fig. 7f). These data reveal that depletion of postnatal Nestin+ perivascular cells caused disruptions in ovarian development and demonstrate that perivascular cells are essential for the breakdown of germline cysts and subsequent follicular formation.

## Discussion

In the adult human ovary, de novo angiogenesis composed of endothelial and perivascular cells occurs in the theca layer of follicles in the early process of follicular growth^39^. Additionally, around 10% of human ovarian cortex cells were annotated as perivascular cells in a recent single-cell study^18^; however, the functional roles of vasculature and perivascular cells in ovarian function are not fully understood. Thus, addressing the roles of perivascular cells is essential for understanding the underlying mechanisms underlying ovarian development. In this study, we identify Nestin+ cells as a source of cortical granulosa cells that are maintained in a perivascular niche via vascular-mediated Notch signaling. Strikingly, depletion of Nestin+ cells in postnatal ovaries impaired the breakdown of germline cysts and disrupted follicular development, likely due to insufficient pre-granulosa cells to surround individual oocytes. Our findings in this study, thus, reveal a critical functional role of vasculature and perivascular cells in supporting folliculogenesis.

Nestin, originally characterized as a marker of neural stem cells, is expressed in stem/progenitor cells in diverse tissues and organs^14, 25, 40^. The role of *Nestin* and *Nestin*-derived cells has been examined in several biological settings, although mostly in a neural context. The *Nestin*^−*/*−^ mutant mouse model exhibits embryonic lethality after E8.5 due to massive apoptosis of neural tube cells^41^. In an adult *Nestin*-CreER;*Rosa*-*lacZ*-DTA mouse model, ablating Nestin+ cell reduced perivascular mesenchymal stromal cells, but did not affect mature bone-marrow-derived Schwann cells^42^. In the reproductive system, Nestin+ cells are putative progenitors for fetal and adult Leydig cells in the testis^14, 25^. In ovaries, Nestin induced by luteinizing hormone (LH) surge via VEGF signal promotes angiogenesis in the follicle^43^, but Nestin+ cells’ specific roles in the ovary were unclear, representing a major outstanding question in the field. Here, we demonstrate that perivascular Nestin+ cells are bona fide cortical granulosa cell progenitors, thus uncovering a new cellular source for granulosa cells.

A common belief is that Sertoli cells and granulosa cells, supporting cells in the testis and ovary, respectively, arise from a common cell lineage in the gonadal primordium^44, 45^. Our lineage-tracing experiments here showed that ovarian Nestin+ cells gave rise to granulosa cells. However, we previously reported that testicular Nestin+ cells do not give rise to Sertoli cells^14^. We also previously showed that testicular Nestin+ cells are a WT1-positive progenitor population with a mesonephric origin^14^, but in the female gonad, here we found that Nestin+ cells are a NR5A1-positive gonadal mesenchymal population at early fetal stages. These results indicate that granulosa cells, in particular ones that arise later in development, and Sertoli cells do not strictly arise from a common precursor. In addition, in contrast to testicular Nestin+ cells that possess self-renewal capacity to expand their numbers^14, 24^, in the fetal ovary new Nestin+ progenitors are instead continuously induced or recruited following blood vessels’ ongoing formation. Moreover, whereas Sertoli cell precursors seem to be mostly derived during a tight time window from a single cellular source (i.e., coelomic epithelium)^46^, granulosa cells in the ovary appear to have multiple origins and a wide window in which they are specified. These differences in gonadal supporting cell specification may be due to the inherent distinctions between spermatogenesis and oogenesis and the kinetics of germline-soma interactions in the testis versus ovary.

Our lineage-tracing assays showed different behaviors of Nestin+ cells at different stages of ovary development. At E15.5, likely the developmental stage during which cortical granulosa cell progenitors begin to arise, Nestin+ progenitors were only adjacent to blood vessels. Instead, most Nestin+ granulosa progenitors emerged between E18.5 and P2, the time when germline cyst breakdown and formation of primordial follicles take place. This time window is likely when the maximum number of granulosa cells is needed to envelop individual oocytes completely and efficiently. In contrast, fewer granulosa cells arising from Nestin+ progenitors at P4 could be observed. Since at P4 the formation of primordial follicles is almost complete^47^ and there is no need for further granulosa cell specification, most Nestin+ cells devoted to a granulosa cell fate have already differentiated and no longer express Nestin. Our long-term lineage tracing results showed that Nestin+ cells present at P4 displayed multipotency and later in development differentiated into multiple interstitial cell types, such as theca cells, pericytes, and interstitial steroidogenic cells, which are required after the establishment of primordial follicles is complete and the ovarian environment needs to be primed for follicular progression. This later *in vivo* multipotency is also a feature of testicular Nestin+ cells^14^ and reveals a similarity between Nestin+ cells in the two sexes.

The vasculature is intimately associated with the maintenance and differentiation of Nestin+ progenitors in the fetal testis^14^, but vasculature’s functional role, if any, in the ovary was as of yet unclear. Here we explored the mechanism of vascular-mesenchymal crosstalk in the developing ovary. We found that vascular disruption reduced the number of Nestin+ progenitors, resulting in precocious or ectopic differentiation of Nestin+ cells into FOXL2-expressing cortical pre-granulosa cells; our analysis of *Foxl2* mRNA levels indicated that there was a significant increase in differentiated pre-granulosa cells when vasculature was disrupted around E18.5, supporting this idea. As a likely result, the reduction of progenitors in fetal ovaries would subsequently lead to a deficit of pre-granulosa cells, thus disrupting the kinetics and/or efficiency of folliculogenesis. Our *Nestin*-CreER;*Rosa-*DTA results support the idea that ablation of postnatal Nestin+ cells impaired the breakdown of germline cysts and subsequent folliculogenesis. Similarly, knockout of *Foxl2* impairs germline cyst breakdown by disrupting the differentiation of granulosa cells and the basal lamina surrounding follicles^48^. Therefore, depletion of Nestin+ cells in postnatal ovaries likely resulted in a reduction of somatic cells surrounding the cysts, and they were subsequently unable to enclose individual oocytes, thus uncovering the importance of Nestin+ progenitor cells during ovary development (Fig. 7h).

Notch signaling frequently plays a crucial role in progenitor cells during embryogenesis by influencing cell proliferation, differentiation, and apoptosis^49^. During ovarian organogenesis, Notch signaling directs the early stages of germline cyst breakdown and primordial follicle formation and maintenance^50, 51^, but how and where Notch was specifically activated was unclear. The early stages of follicle assembly appear to be the most sensitive to Notch signaling, as there were no significant differences in later-stage follicle populations after blockade of Notch activity^50^. Additionally, Notch signaling in fetal ovaries promotes the formation of primordial follicles via the regulation of oocyte survival and proliferation of pre-granulosa cells^52^. Since most research into ovarian Notch signaling has focused on oocyte-granulosa cell interactions, Notch’s role in the ovarian interstitium was unclear. Here we found that in *CBF*:H2B-Venus Notch reporter mice, Venus expression was increased at E18.5 (onset of the breakdown of germline cysts), consistent with the studies of Notch family gene expression in the neonatal mouse ovary^50^. However, Notch activity in the fetal and perinatal ovary was detected within perivascular mesenchymal cells, implicating vasculature as an essential mediator of ovarian Notch signaling.

While vasculature is required for interstitial Notch signaling in the fetal testis, whereby endothelial cells provide Notch ligand to perivascular Nestin+ cells^14^, the importance of vasculature for Notch signaling in the developing ovary was unknown. The depletion of vasculature in *CBF*:H2B-Venus ovaries *ex vivo* showed that gonadal and medullary Notch activity, as well as Nestin expression, depends on the vasculature. However, the vasculature is also impaired by blocking Notch signaling at later stages, indicating that there are mutual or reciprocal effects of Notch on endothelial and mesenchymal cells: in early stages, activity of Notch depends on the vasculature, while later on, activation of Notch signaling might contribute to vasculature formation. Similar to vascular disruption, blockade of Notch signaling induced the ectopic or precocious differentiation of Nestin+ cells into cortical pre-granulosa cells and reduced medullary perivascular undifferentiated Nestin+ cells at E18.5, further confirming vasculature as a critical regulator of ovarian Notch activity.

Our study reveals a new precursor of granulosa cells, perivascular Nestin+ cells, which are regulated by blood vessels via Notch signaling during ovary organogenesis. Reduction of vascular-dependent Notch activity induced differentiation of Nestin+ progenitors into cortical pre-granulosa cells in fetal ovaries, disturbing the formation of primordial follicles and resulting in polyovular follicles. *In vivo* cell ablation revealed a crucial role for Nestin+ progenitors in the differentiation of pre-granulosa cells, ensuring the timely and efficient formation of primordial follicles. Our data reveal how blood vessels mediate a balance between maintenance and differentiation of ovarian progenitors and their importance for folliculogenesis. These findings provide new insights into ovarian organogenesis which may have implications for our understanding of the etiology of female gonad dysgenesis and infertility.

## Methods

### Mice

CD-1 mice (Charles River) were used for wild-type expression and *ex vivo* organ culture studies. Cre-responsive *Rosa*-Tomato reporter mice (B6.Cg-*Gt(ROSA)26Sor^tm^*^14^*^(CAG-tdTomato)Hze^*; JAX stock #007914), *Rosa*-eGFP-DTA mice (*Gt(ROSA)26Sor^tm^*^1^*^(DTA)Jpmb^*; JAX stock #006331), and *CBF*:H2B-Venus mice (Tg(Cp-HIST1H2BB/Venus)47Hadj; JAX stock #020942), were obtained from The Jackson Laboratory. *Nestin*-CreER mice (Tg(Nes-cre/ERT2,-ALPP)1Sbk)^14, 53^, on a mixed B6/FVB/CD-1 genetic background, were obtained from Dr. Masato Nakafuku (Division of Developmental Biology, Cincinnati Children’s Hospital Medical Center). *Rosa*-NICD mice (*Gt(ROSA)26Sor^tm^*^1^*^(Notch1)Dam^*)^54^ were provided by Dr. Joo-Seop Park (Divisions of Pediatric Urology and Developmental Biology, Cincinnati Children’s Hospital Medical Center). Mice were housed in accordance with National Institutes of Health guidelines, and experimental protocols were approved by the Institutional Animal Care and Use Committee (IACUC) of Cincinnati Children’s Hospital Medical Center (animal experimental protocol number IACUC2018-0027).

### Immunofluorescence

Tissues were fixed in 4% paraformaldehyde (PFA) with 0.1% Triton X-100 overnight at 4 °C. Whole-mount immunofluorescence was performed on gonads at stages E11.5 and E12.5. After several washes in PBS + 0.1% Triton X-100 (PBTx), samples were incubated in blocking solution (PBTx + 10% FBS +3% bovine serum albumin [BSA]) for 1 hour at room temperature. Gonads were incubated with primary antibodies (diluted in blocking solution) on a rocker overnight at 4 °C. After several washes in PBTx, fluorescent secondary antibodies were applied for 4 hours rocking at room temperature. After several washes in PBTx, samples were mounted on slides in Fluoromount-G (SouthernBiotech).

For fetal ovaries at stages E15.5 and older, cryosections were performed. Dissections and fixation were performed as above. After several washes in PBTx, samples were processed through a sucrose:PBS gradient (10, 15, 20% sucrose) before an overnight incubation in a 1:1 mixture of 20% sucrose and OCT medium (Sakura) rocking at 4 °C. Samples were embedded in OCT medium and stored at −80 °C. Cryosections (12 µm thickness) from control and experimental groups were processed onto the same slide. Cryosections were then stained and mounted as above, except secondary antibodies were applied for only 1 hour.

Primary antibodies and dilutions used in this study are listed in Supplementary Table 1. *Rosa*-Tomato and *CBF*:H2B-Venus expression were detected via endogenous fluorescence. Alexa-488-, Alexa-555-, and Alexa-647-conjugated secondary antibodies (Molecular Probes/Life Technologies/Thermo Fisher) were used at 1:500. Nuclear staining was performed using 2 μg/mL Hoechst 33342 (#H1399, Thermo Fisher), and Hoechst staining is labeled as “Nuclei” in all figures. Pictures were taken on a Nikon Eclipse TE2000 microscope (Nikon, Tokyo, Japan) equipped with an Opti-Grid structured illumination imaging system running Volocity software (PerkinElmer, Waltham, MA, USA) or on a Nikon A1 Inverted Confocal Microscope (Nikon, Tokyo, Japan). At least three independent experiments were performed and within each experiment multiple samples (*n*≥2) were used.

### Lineage tracing of Nestin+ cells

For *in vivo Nestin*-CreER experiments, CreER-positive males were crossed with *Rosa*-Tomato females. Pregnant females were injected intraperitoneally at E12.5, E15.5, E18.5, P2, or P4 with 75 μg/g 4-hydroxytamoxifen (4-OHT; Sigma-Aldrich #H6278) dissolved in corn oil. For E12.5 injection experiments, an additional progesterone (37.5 μg/g; Sigma-Aldrich #P0130) injection was given subcutaneously at E14.5 to help maintain pregnancy further after 4-OHT administration^14^. Pregnant females were euthanized at the required stage, and ovaries from embryos or pups were dissected and processed for immunofluorescence or organ culture. For lineage tracing into adulthood, ovaries from *Nestin*-CreER; *Rosa*-Tomato females were analyzed at 30 days (P30) and 60 days (P60) old (*n*=4 females for each stage).

### Ablation of Nestin+ cells

To specifically deplete Nestin+ cells from the ovary, the Cre-responsive *Rosa*-eGFP-DTA system^37^ was used. *Nestin*-CreER;*Rosa-*Tomato homozygous males were crossed to *Rosa*-eGFP-DTA heterozygous females. Pups were intragastrically injected with 4-OHT (0.05 mg/each pup, dissolved in corn oil) at P2 and P4 to target perivascular Nestin+ cells. Diphtheria toxin is expressed only in CreER-active cells, thus inducing apoptosis only in perivascular cells. *Rosa*-Tomato reporter expression was used in the genetic background to assess efficiency of depletion of Nestin+ cells. Control *Nestin*-CreER;*Rosa-*Tomato and cell-ablated *Nestin*-CreER;*Rosa-*Tomato;*Rosa*-eGFP-DTA ovaries (*n*=4 females for each genotype and stage) were harvested and analyzed at P7 and P21.

### MitoTracker coelomic epithelial labeling

E11.5 and E12.5 CD-1 gonads were incubated in 1 mM MitoTracker Orange CMTMRos (Thermo Fisher Scientific/Invitrogen #M7510) in PBS for 30 minutes at 37 °C to label the surface epithelium. Following incubation with MitoTracker, gonads were washed several times in culture media. Gonads were then cultured for 48 hours in 1.5% agar wells and processed for whole-mount immunofluorescence as described above. As a control, several gonads in each experiment were removed and fixed immediately after labeling and washing with culture media to ensure that only the surface coelomic epithelial cells were initially labeled. At least three independent experiments were performed and within each experiment multiple samples (*n*≥2) were used.

### Flow cytometry

*Nestin*-CreER-positive males were crossed with *Rosa*-Tomato females. Pregnant females were injected intraperitoneally at E12.5 with 75 μg/g 4-hydroxytamoxifen (4-OHT; Sigma-Aldrich #H6278) dissolved in corn oil, and an additional progesterone (37.5 μg/g) injection was given subcutaneously at E14.5 to help maintain pregnancy further after 4-OHT administration^14^. E18.5 fetal ovaries were dissected and pooled in a 1.5-ml microcentrifuge tube. Similarly, E18.5 fetal ovaries from control littermates were used for negative and unstained controls. 0.05% Trypsin-EDTA (Thermo Fisher Scientific/Gibco #25300054) was added to the pooled gonads (50 μl/gonad) and placed at 37°C for 6 minutes, with periodic tapping at 2-minute intervals. Digestion was stopped by adding an equal volume of culture media supplemented with 10% FBS. Trypsin mixture was removed carefully without disturbing the tissue on the bottom of the tubes. 1 ml of 10% FBS solution (in 4°C PBS) was added to each tube. Using a 1000p pipette set at 300μl, samples were gently pipetted until the tissue was completely dissociated. After cellular dissociation, the cells were briefly centrifuged to spin down the cell pellet (600 x *g* for 5 minutes at 4°C) and resuspended in PBS with 10% FBS and kept on ice until flow cytometric analysis.

Cell suspensions from whole embryonic female gonads were filtered through a 35-micron sieve (BD Biosciences). Cells were analyzed using a BD LSRFortessa running BD FACSDiva v8.0 software to detect endogenous fluorescence signal in tdTomato+ (RFP+) cells versus tdTomato-negative (RFP-negative) cells from *Nestin*-CreER;*Rosa*-Tomato ovaries. Analysis and quantification were performed using FlowJo v10 software. Ovaries from *n*=3 independent embryos were used for each stage.

### Histological analysis of ovaries and follicle quantification

Entire ovaries were serially sectioned and stained for counting. Paraffin-embedded ovary sections (5 μm) were deparaffinized and rehydrated through a series of xylene and ethanol. Sections were stained with Hematoxylin (Denville) and incubated for 5 minutes. Follicles were counted in every 5th section through the entire ovary from each mouse as described previously^55^. All follicle stages were categorized according to morphological criteria^38^. Briefly, primordial follicles contained an oocyte surrounded by squamous cells; primary follicles were classified as oocytes surrounded by a single layer of cuboidal granulosa cells; secondary follicles were regarded as oocytes surrounded by more than one layer of granulosa cells; antral follicles showed a visible cavity and large antral follicle contained a large antral cavity. For secondary to large antral follicles, we only counted follicles with a visible nucleus to avoid double counting. The total number of follicles was defined as the sum of the number of all stages of follicles in one ovary. For counting follicles with Tomato+ granulosa cells, only primary, secondary, and antral follicles were counted. Control *Nestin*-CreER;*Rosa-*Tomato and cell-ablated *Nestin*-CreER;*Rosa-*Tomato;*Rosa*-eGFP-DTA ovaries (*n*=4 for each genotype and stage) were harvested and analyzed at P7 and P21.

### Ex vivo whole-organ ovarian culture

E12.5 and E15.5 gonad-mesonephros complexes or E18.5 and P2 isolated ovaries were dissected from CD-1, *Nestin*-CreER;*Rosa*-Tomato, or *CBF*:H2B-Venus embryos or pups. Ovaries were cultured using an agar culture method as previously described^56^. Briefly, organs were positioned on 1.5% agarose gel placed in petri dishes with complete medium (DMEM/10% fetal bovine serum [FBS]/1% penicillin–streptomycin) at 37°C and 5% CO_2_. Medium was changed every 24 hours. 1.8 μg/mL VEGF Receptor Tyrosine Kinase Inhibitor II (VEGFR-TKI II; Calbiochem/EMD Millipore #676481-5MG) dissolved in DMSO was used to disrupt blood vessels^57^ and 100 μM γ-secretase inhibitor IX (DAPT; Calbiochem/EMD Millipore #565770-5MG) dissolved in DMSO was used to block γ-secretase activity^14^. Following culture for 72 hours, E12.5 gonad-mesonephros complexes, fetal ovaries, and postnatal ovaries were either fixed for immunofluorescence or the gonads (separated from mesonephros) were collected for RNA extraction and qRT-PCR analyses.

In *Nestin-*CreER;*Rosa-*Tomato *ex vivo* culture experiments, 4-OHT was dissolved in ethanol and added at 0.02 mg/mL in the culture media to induce CreER activity. To eliminate the possibility that vascular disruption or Notch inhibition by DAPT could hinder the ability of Tomato+ cells to be specified initially, 4-OHT alone was added to the culture media for the first 24 hours of culture, after which 4-OHT was removed and VEGFR-TKI II or DAPT was added to the culture media for the remaining additional 72 hours of culture. Preliminary experiments revealed that Tomato reporter expression was already robustly activated after the first 24 hours of culture (not shown). At least three independent experiments were performed and within each experiment multiple gonads (*n*≥2) were used.

### RNA extraction and quantitative real-time PCR (qRT-PCR)

Total RNA was extracted from gonads (separated from the mesonephros) using a standard TRIzol (Invitrogen/Thermo Fisher) and isopropanol precipitation protocol. cDNA synthesis was performed with 500 ng of total RNA, using an iScript cDNA synthesis kit (Bio-Rad). The cDNA was then subjected to qRT-PCR using the Fast SYBR Green Master Mix (Applied Biosystems/Thermo Fisher) on the StepOnePlus Real-Time PCR System (Applied Biosystems/Thermo Fisher). The following parameters were used: 95 °C for 20 s, followed by 40 cycles of 95 °C for 3 s and 60 °C for 30 s. Primer specificity for a single amplicon was verified by melt curve analysis or agarose gel electrophoresis. Primers used for qRT-PCR are listed in Supplementary Table 2. All reactions were run in triplicate. *Gapdh* was used as an internal normalization gene. Data from qRT-PCR was calculated relative to controls using the ΔΔCt method. Results were shown as mean ± SD. A two-tailed Student t-test was performed to calculate *P* values, in which *P* < 0.05 was considered statistically significant. Statistical analyses were performed using Prism version 5.0 (GraphPad). At least two independent experiments were performed and within each experiment multiple fetal or postnatal ovaries (*n*≥3) were used.

## Data availability

The data that support the findings of this study are available from the authors on reasonable request.

## Acknowledgements

We thank Drs. Masato Nakafuku and Joo-Seop Park for mice and Dr. Dagmar Wilhelm for anti-FOXL2 antibody. We acknowledge BioRender software (biorender.com) for creation of figures. This work was supported by Lalor Foundation (postdoctoral fellowship to S.L.) and by National Institutes of Health (grants R35GM119458 and R01HD094698 to T.D.).

## Author contributions

S.L. conducted experiments, performed data analyses, and co-wrote and edited the manuscript. B.B. conducted experiments and edited the manuscript. T.D. supervised the project and co-wrote and edited the manuscript.

## Competing interests

The authors declare no competing interests.

## Notes

### Competing Interest Statement

The authors have declared no competing interest.

