## Supplemental Data for "Perivascular cells support folliculogenesis in the developing ovary"

**Supplementary Figure Legends**

**Supplementary Fig. 1. Gonadal NR5A1+ cells are a source of Nestin+ cells in the fetal ovary.**

(a-f) Immunofluorescence images of E11.5 (a,b) or E12.5 (c-f) fetal ovaries labeled with Mitotracker and cultured *ex vivo*. (a) E11.5 gonad immediately after initial 30-minute incubation with MitoTracker dye, showing labeling limited to coelomic surface epithelial cells. (b) After 48-hour culture, coelomic epithelial cells labeled by MitoTracker migrated into the gonad. MitoTracker label did not overlap with Nestin. (c) E12.5 gonad immediately after initial 30-minute incubation with MitoTracker dye, showing labeling limited to surface coelomic epithelial cells. (d-f) After 48-hour culture, epithelial cells labeled by MitoTracker migrated into the gonad. MitoTracker label did not overlap with Nestin (e). MitoTracker-labeled epithelial cells that remained on the coelomic surface co-expressed Nestin (f, arrowheads). (g,h) Immunofluorescence images of E13.5 *Nestin*-CreER;*Rosa*-Tomato fetal ovaries exposed to 4-hydroxytamoxifen (4-OHT) at E12.5. Nestin+ cells are NR5A1-positive (g) but WT1-negative (h). b', g', and h' are higher-magnification images of the boxed regions in b, g, and h. e and f are higher-magnification images of the corresponding boxed regions in d. Dashed lines indicate gonad-mesonephros border. Scale bars: 50 μm.

**Supplementary Fig. 2. Initial Tomato labeling in Nestin+ cell lineage tracing marks perivascular cells in fetal and postnatal ovaries.**

(a-d) Immunofluorescence images of *Nestin*-CreER;*Rosa*-Tomato fetal and postnatal ovaries exposed to 4-OHT at E15.5 (a), E18.5 (b), P2 (c), and P4 (d). Images were taken 24 hours after

4-OHT injection, demonstrating initial Tomato reporter labeling. In E15.5-injected samples (a), Tomato-labeled cells were perivascular and mutually exclusive of FOXL2 expression. While at all stages the vast majority of Tomato-labeled cells were FOXL2-negative, occasional Tomato-positive cells co-expressed FOXL2 (b-d, arrowheads). Scale bars: 50 μm.

**Supplementary Fig. 3. Perivascular Nestin+ progenitors are multipotent and give rise to multiple ovarian cell types.**

(a-x) Long-term lineage-tracing experiments for Nestin+ cells in *Nestin*-CreER;*Rosa*-Tomato juvenile (P30) and adult (P60) ovaries, exposed to 4-OHT at E15.5 (a-f), E18.5 (g-l), P2 (m-r), or P4 (s-x). P30 and P60 ovaries were examined for Tomato, steroidogenic cell marker HSD3B1, smooth-muscle cell marker ACTA2 (also called αSMA), and pericyte marker CSPG4 (also called NG2). HSD3B1 was observed not only in the theca cell layer but also in interstitial cells. ACTA2 was expressed in the theca layer. CSPG4 was expressed in both the theca layer and in pericytes. Arrowheads indicate Tomato+ cells of interest labeled by each antibody. Scale bars: 50 μm.

**Supplementary Fig. 4. Pattern and timing of active Notch signaling in fetal and postnatal ovaries.**

(a) Images of E15.5, E18.5, P2, and P4 ovaries from *CBF*:H2B-Venus mice (active Notch signaling reporter). Few ICAM2-labeled endothelial cells (white arrows) and Nestin+ cells adjacent to the vasculature (yellow arrows) were Venus+ at E15.5. At E18.5, the number of Venus+ cells increased, were down-regulated at P2, and by P4 few Venus+ cells were detected in the ovary. (b-e) qRT-PCR analyses showing fold change in Notch target gene (*Hes1* (b), *Hes5* (c), *Hey1* (d), *Heyl* (e)) mRNA levels in ovaries at various developmental stages. Different letters indicate statistically different values ($P<0.05$; two-tailed Student's t-test). qRT-PCR values in b-e are presented as mean ± SD. (f) Image of P2 *CBF*:H2B-Venus ovary. Perivascular Venus+ cells did not express FOXL2 (blue arrowhead), while cortical Venus+ cells co-expressed FOXL2 (white arrowheads). Scale bars: 50 μm.

**Supplementary Fig. 5. Active Notch signaling induces expression of Nestin in perivascular cells.**

(a) qRT-PCR analyses showing fold change in *Cdh5* and *Nestin* mRNA levels after blockade of Notch signaling via DAPT treatment at E18.5 for 12, 18, or 24 hours. *, $P<0.05$, **, $P<0.01$; two-tailed Student's t-test. qRT-PCR values in A are presented as mean ± SD. (b,c) Immunofluorescence images of P1 *Nestin*-CreER; *Rosa*-Tomato; *Rosa*-NICD fetal ovaries exposed to 4-OHT at E18.5, showing that perivascular Nestin-derived Tomato+ cells were increased in NICD+ ovaries as compared to controls; however, Tomato+ cells did not co-localize with MKI67 (Ki67) or phospho-histone H3 (pHH3). Scale bars: 50 μm.

**Supplementary Fig. 6. *In vivo* ablation of Nestin+ cells in the postnatal ovary is highly efficient.**

(a,b) Immunofluorescence images of P7 (a) and P21 (b) control (*Nestin*-CreER;*Rosa*-Tomato) and DTA+ (*Nestin*-CreER;*Rosa*-Tomato;*Rosa*-eGFP-DTA) postnatal ovaries exposed to 4-OHT at P2 and P4. DTA+ ovaries exhibited a complete ablation of Tomato+ cells. In controls,

Tomato+ cells co-expressed the granulosa cell marker FOXL2 in both P7 and P21 ovaries. GCNA (TRA98) labels germ cells. Scale bars: 50 µm.

**Supplementary Table 1. Primary antibodies used for immunofluorescence.**

| Primary Antibody | Dilution | Source/Reference |
|---|---|---|
| Rabbit anti-ACTA2 (αSMA) | 1:500 | Abcam #ab5694 |
| Rabbit anti-CSPG4 (NG2) | 1:500 | Millipore-Sigma #AB5320 |
| Rabbit anti-FOXL2 | 1:500 | D. Wilhelm (Polanco, et al., 2010) |
| Rat anti-GCNA (TRA98) | 1:1,000 | Abcam #ab82527 |
| Rabbit anti-HSD3B1 | 1:500 | Cosmo Bio #KAL-KO607 |
| Rat anti-ICAM2 (CD102) | 1:1,000 | Bio-Rad #MCA2295 |
| Rabbit anti-MKI67 (Ki67) | 1:500 | GeneTex #GTX16667 |
| Rabbit anti-Nestin | 1:1,000 | BioLegend #PRB-315C |
| Rat anti-NR5A1 | 1:200 | Cosmo Bio #KAL-KO610 |
| Rat anti-PECAM1 | 1:250 | BD Pharmingen #553370 |
| Rabbit anti-phospho-histone H3 (Ser10) | 1:500 | Millipore-Sigma #06-570 |
| Mouse anti-WT1 (F-6) | 1:100 | Santa Cruz #sc-7385 |

**Supplementary Table 2. Sequences of primers used for qRT-PCR analyses.**

| **Gene name** | **Sequence (5' to 3')** |
|---|---|
| *Cdh5* forward | TCCTCTGCATCCTCACTATCACA |
| *Cdh5* reverse | GTAAGTGACCAACTGCTCGTGAAT |
| *Nestin* forward | GCTGGAACAGAGATTGGAAGG |
| *Nestin* reverse | CCAGGATCTGAGCGATCTGAC |
| *Hes1* forward | ATAGCTCCCGGCATTCCAAG |
| *Hes1* reverse | GCGCGGTATTTCCCCAACA |
| *Hes5* forward | AGTCCCAAGGAGAAAAACCGA |
| *Hes5* reverse | GCTGTGTTTCAGGTAGCTGAC |
| *Hey1* forward | GCGCGGACGAGAATGGAAA |
| *Hey1* reverse | TCAGGTGATCCACAGTCATCTG |
| *Heyl* forward | CAGCCCTTCGCAGATGCAA |
| *Heyl* reverse | CCAATCGTCGCAATTCAGAAAG |
| *Foxl2* forward | GCTACCCCGAGCCCGAAGAC |
| *Foxl2* reverse | GTGTTGTCCCGCCTCCCTTG |
| *Gapdh* forward | AGGTCGGTGTGAACGGATTTG |
| *Gapdh* reverse | TGTAGACCATGTAGTTGAGGTCA |

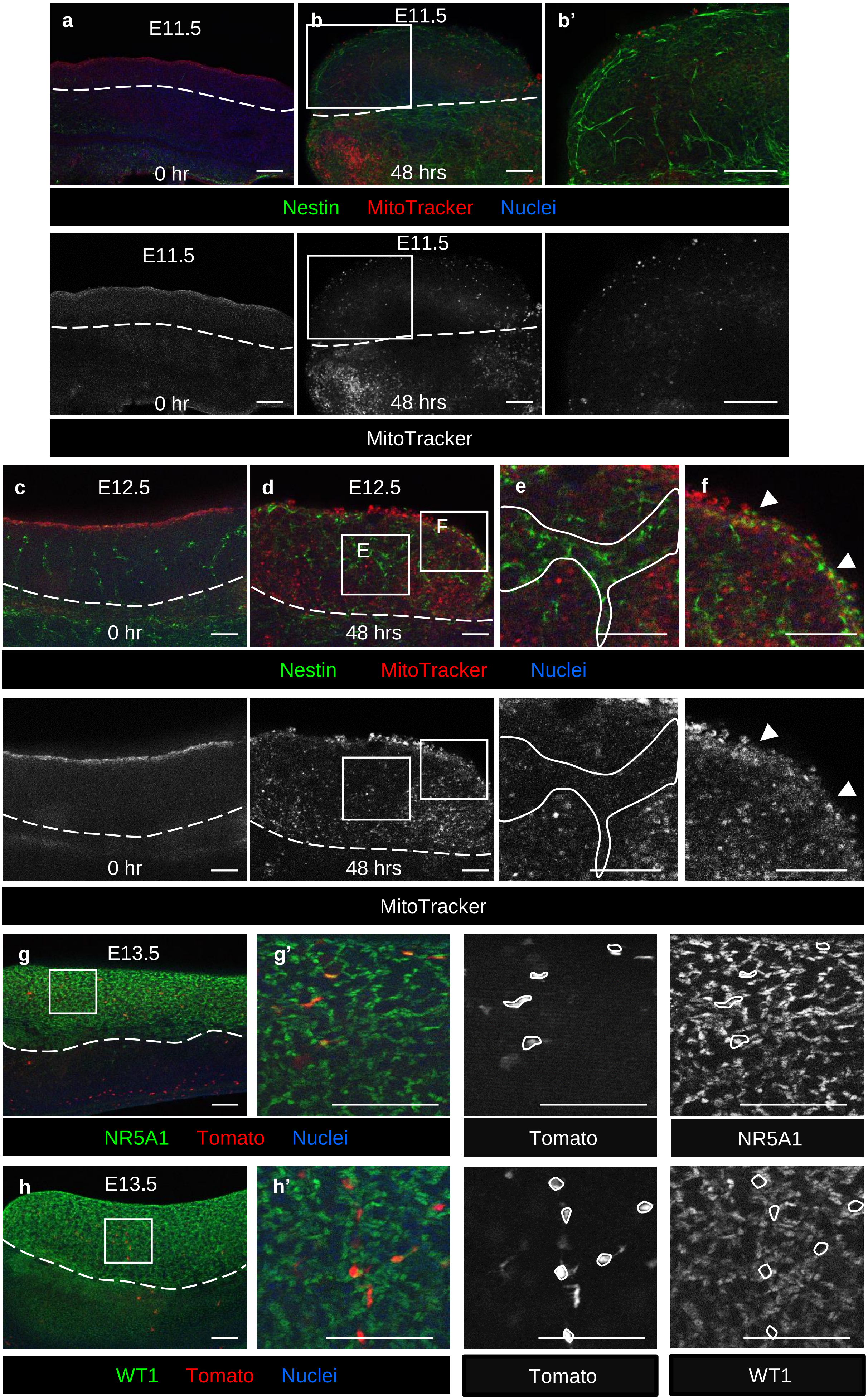


**Supplemental Fig. 1**

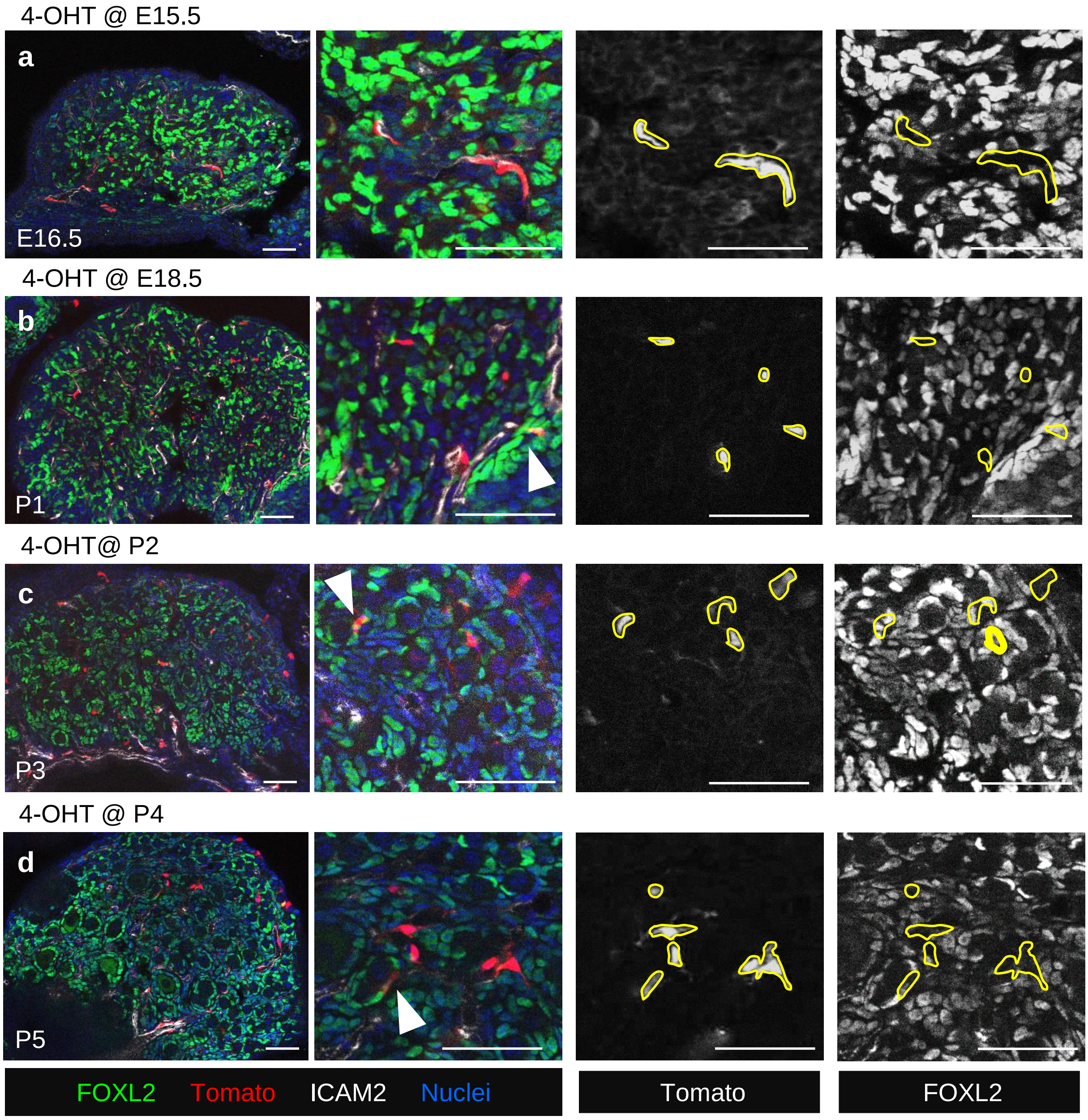


**Supplemental Fig. 2**

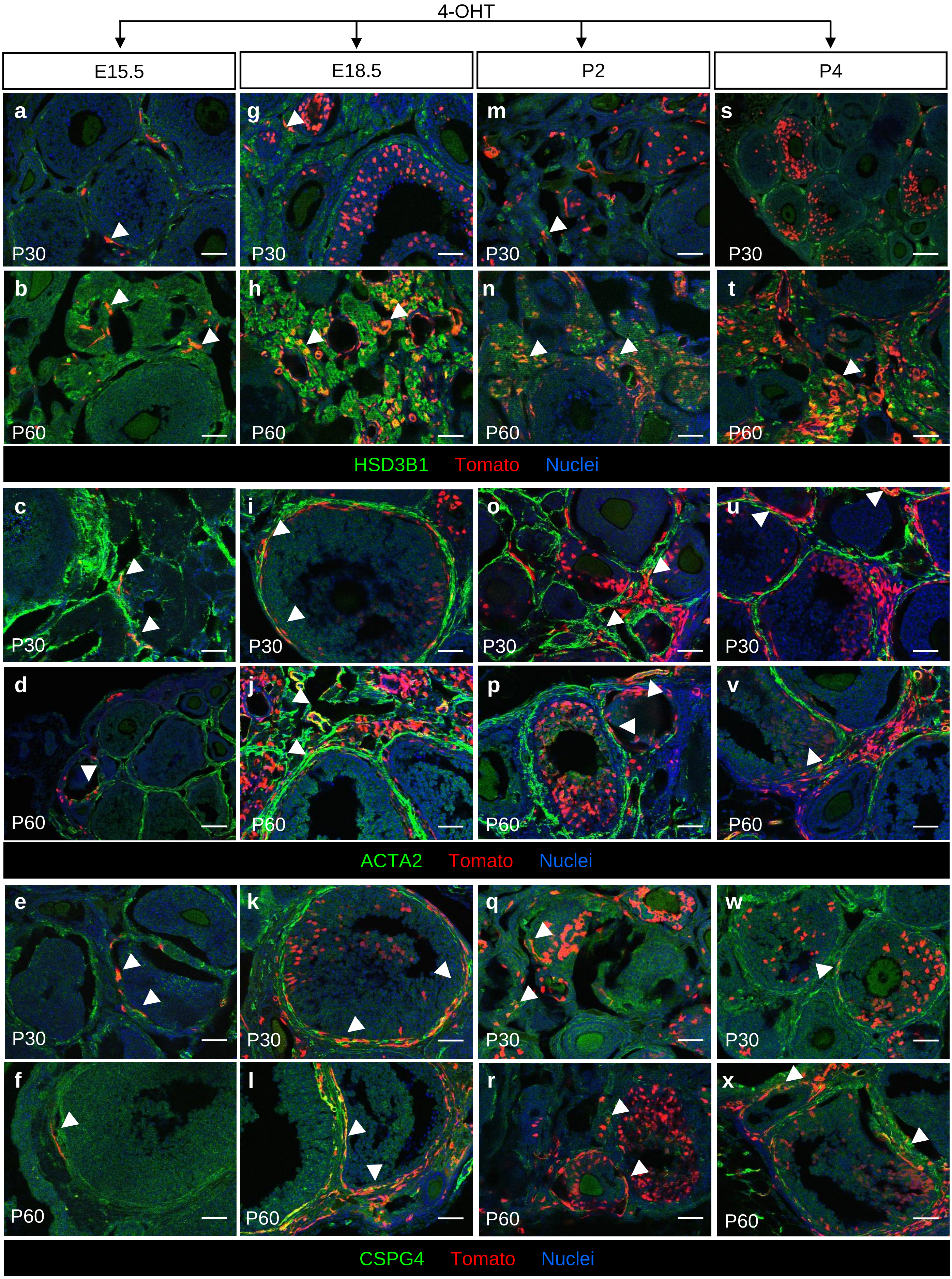


**Supplemental Fig. 3**

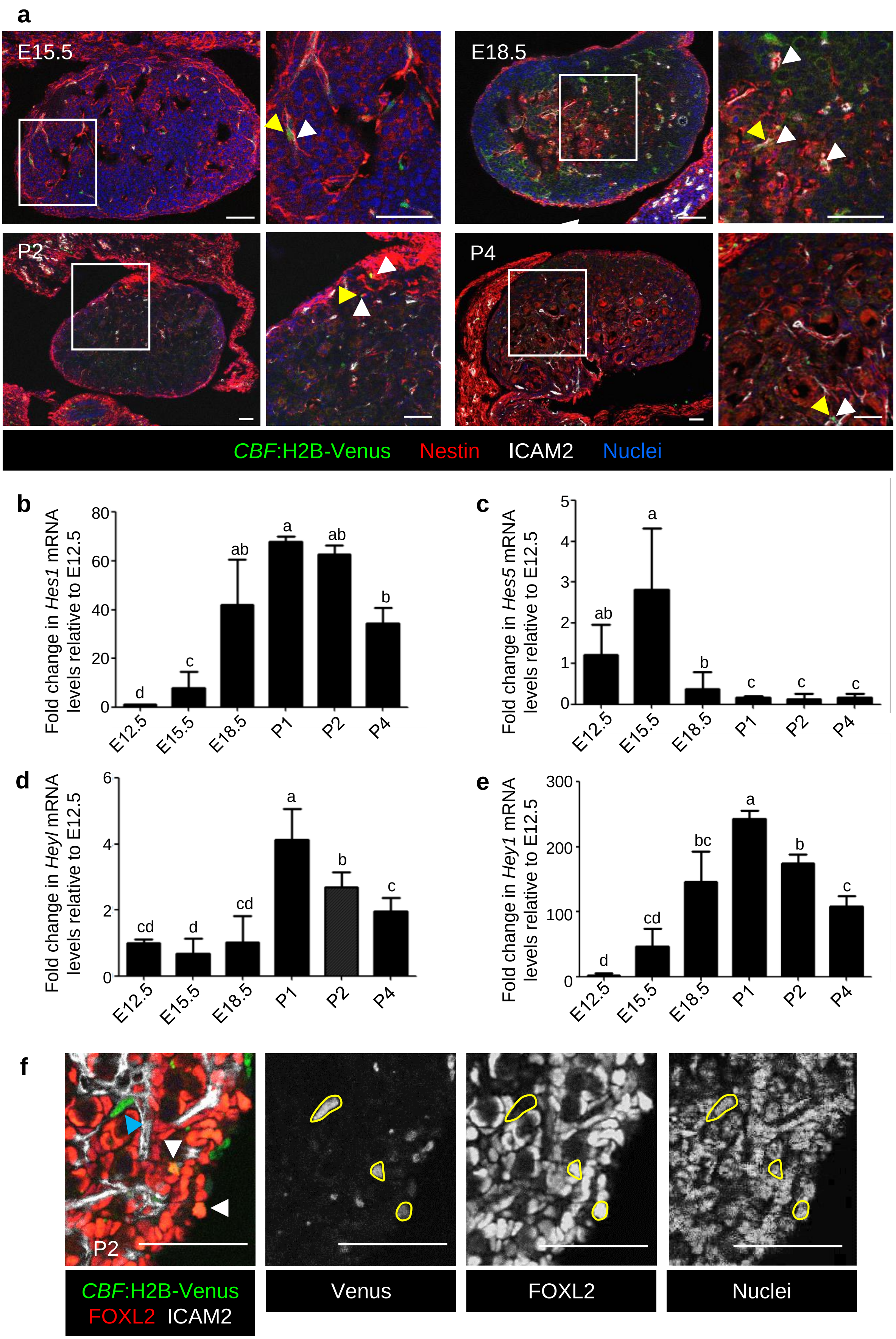


**Supplemental Fig. 4**

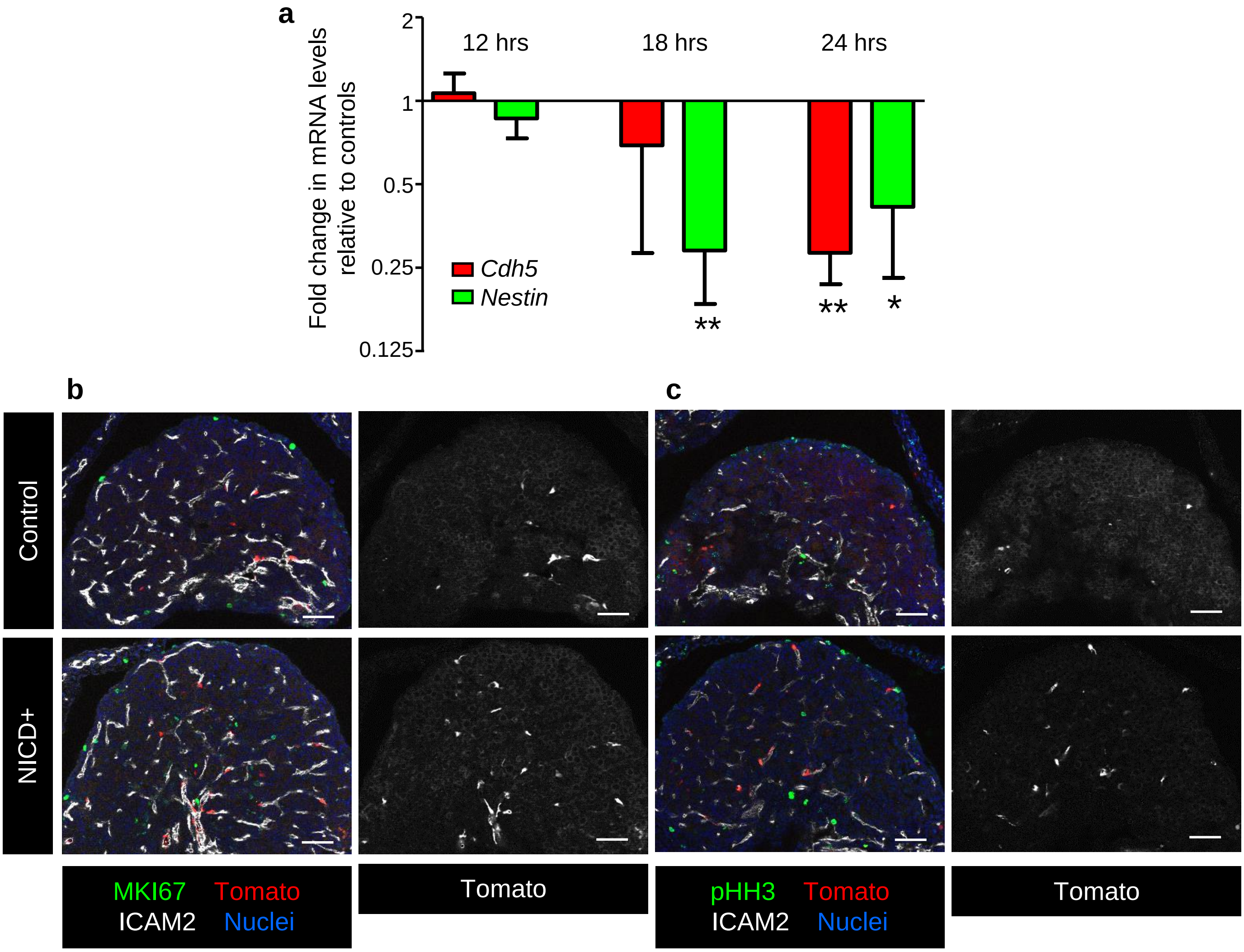


**Supplemental Fig. 5**

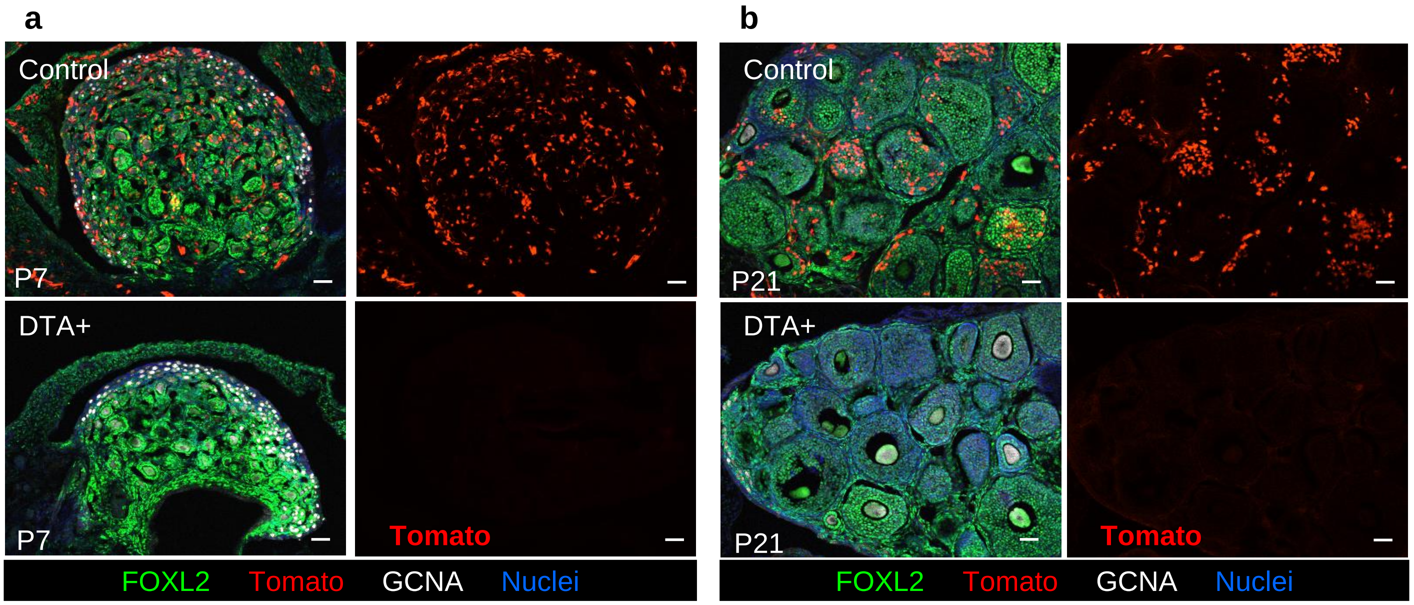


**Supplemental Fig. 6**
